# Evaluation of harmonization methods to mitigate assay and cohort effects in plasma p-tau217

**DOI:** 10.64898/2026.07.24.740589

**Authors:** Vivian Z. Zhang, Pamela C.L. Ferreira, Yiwen Dong, Davneet Minhas, Guilherme Povala, Bruna Bellaver, Tharick A. Pascoal, Xuemei Zeng, Thomas K. Karikari, Ann D. Cohen, Rebecca A. Deek, Qiong Wu, Dana L. Tudorascu

**Affiliations:** Department of Biostatistics and Health Data Science, University of Pittsburgh School of Public Health; Department of Psychiatry, University of Pittsburgh School of Medicine; Department of Radiology, University of Pittsburgh School of Medicine; Department of Neurology, University of Pittsburgh School of Medicine; Alzheimer’s Disease Research Center, University of Pittsburgh

## Abstract

**INTRODUCTION:** The growing number of assay platforms measuring blood-based biomarkers (BBMs) for Alzheimer’s disease (AD) has introduced challenges in interpretability and comparability across assays. Differences across studies also limit comparability of data. To address these challenges, a systematic evaluation of harmonization methods is needed to support BBM data integration within or across studies.

**METHODS:** Two multisite studies, Alzheimer’s Disease Neuroimaging Initiative (ADNI, n = 219) and Human Connectome Project (HCP, n = 111), were used to evaluate harmonization methods for mitigating assay and cohort effects in plasma p-tau217 measurements. Methods includes various normalization, regression, and standardization approaches, including the recently developed CentiMarker. Assay effects were evaluated using repeated-measures data across assay platforms within each cohort, whereas cohort effects were assessed using pooled ADNI and HCP data. Harmonization performance was evaluated using distributional statistics and downstream modeling of p-tau217.

**RESULTS:** Quantile normalization and quantile mapping methods were most effective for mitigating assay effects, whereas conditional quantile mapping performed best for pooled multi-cohort data. These methods also preserved biological variability. In contrast, simple means adjustment and reference-based z-score standardization were least effective for mitigating assay effects, while simple means adjustment, z-score standardization, and quantile normalization were least effective for mitigating cohort effects. CentiMarker had minimal impact on assay or cohort effects.

**DISCUSSION:** Based on our evaluation, we recommend (conditional) quantile mapping for p-tau217 studies integrating data across multiple assays or cohorts. In contrast, we caution against using CentiMarker and z-score-based methods, as they limit comparability and do not effectively mitigate technical variability.

## 1 Introduction

In recent years, several blood-based biomarkers (BBM) have been developed to detect Alzheimer’s disease (AD) pathology. BBMs offer a scalable and cost-effective approach for assessing AD-related neuropathology, addressing many practical limitations of positron emission tomography (PET) and CSF testing.^1^ As a result, there has been substantial interest in applying BBMs across large epidemiological studies, clinical trials, and primary care settings to aid in disease screening and monitoring of neuropathological changes.^2,3^ Among these BBMs, phosphorylated tau (p-tau), particularly p-tau217, has emerged as a promising candidate. Studies suggest that plasma p-tau217 reflects early amyloid-beta (Aβ) alterations in the brain and demonstrates high diagnostic performance for identifying amyloid plaques and differentiating AD from other neurodegenerative disorders.^4-6^

However, the expanding use of p-tau217 across diverse assay platforms has introduced new challenges. Recent studies have reported substantial variability in quantitative p-tau217 measurements across assay platforms, known as assay (platform) effects.^3,5-9^ Comparative analyses have identified significant proportional bias across all assay pairs and significant systematic bias in most pairwise comparisons^7-9^, indicating that p-tau217 values are not directly interchangeable across assays. This inter-assay variability likely arises from differences in assay technology, protocols, and antibody design^3,9^, thereby limiting comparability between assays. Moreover, inconsistencies across cohort studies exacerbate unwanted variation in the data, known as cohort effects, and limit the comparability of findings across studies.

To address these challenges, harmonization methods^10^ are needed to reduce unwanted technical variability and enable meaningful comparisons within or across studies. Comparative studies of commercially available p-tau217 immunoassays have demonstrated strong correlations between assays^5,7,8^, suggesting that measurements are systematically related and may be combined using appropriate harmonization methods during data preprocessing. One method commonly used to improve comparisons of plasma biomarkers is z-score standardization^11^, which involves using a reference group to convert raw or log-transformed values into z-scores. More recently, researchers proposed the CentiMarker metric to standardize AD biomarker values on a scale from 0 to 100 for monitoring disease progression or treatment response^12^. Additionally, research in omics and cancer microarrays has developed harmonization methods that offer promising, adaptable methods for addressing assay effects in BBM data.^13-16^ With these various harmonization methods, a comprehensive evaluation is needed to assess their ability to mitigate technical sources of variability, including assay and cohort effects.

In this study, we provide a novel application of existing harmonization methods to address technical variability in plasma p-tau217 data measured across multiple assay platforms from two independent cohorts. This study aimed to systematically evaluate these methods to mitigate (a) assay effects in repeated measures of p-tau217 data within two cohorts, and (b) cohort effects in pooled data within shared assay platforms. An effective harmonization method will decrease technical variability while preserving biologically meaningful differences, including mean differences between cognitively impaired (CI) and cognitively unimpaired (CU) groups.

## 2 Methods

We performed two complementary data harmonization analyses: (1) evaluation of assay platform effects within cohorts using their repeated-measures data, and (2) evaluation of cohort effects using pooled data across cohorts from shared assay platforms. Both analyses followed a common procedure: application of harmonization methods, followed by evaluation of technical and biological variability. **Appendix p 1** illustrates the analytical workflows, including the datasets, harmonization methods, and evaluation metrics used in each setting.

### 2.1 Participants

We included 330 participants from two cohorts, each with repeated measures of plasma p-tau217 from multiple assay platforms. The characteristics of each cohort are described below:

#### Cohort 1: ADNI dataset

The dataset includes 219 participants from the Alzheimer’s Disease Neuroimaging Initiative (ADNI).^17^ Each participant had p-tau217 measurements from four assays: ALZpath (Quanterix), Janssen p-tau217+ (Quanterix), Fujirebio Lumipulse, and C2N PrecivityAD2. 172 participants with at least one assay value below the limit of quantification or outside the calibration range were removed, resulting in the final sample size of 219. Across assays, excluded values included 172 from C2N, one from ALZpath, and one from Janssen. Participants were classified as CU if they had a Clinical Dementia Rating (CDR) of 0, and as CI if they had a CDR > 0.

#### Cohort 2: HCP dataset

The dataset includes 111 participants from the Human Connectome Project (HCP) recruited at the University of Pittsburgh. The HCP is a community-based cohort conducted at the University of Pittsburgh as part of the NIA-funded Human Connectome Project initiative.^18^ Each participant had measurements from ALZpath, Janssen, Fujirebio, and the University of Pittsburgh in-house single-molecule array assay.^19^ Diagnostic groups include CU for participants with a clinical classification of normal control, impaired without complaints, or subjective cognitive decline, and CI for those with a clinical classification of amnestic mild cognitive impairment, non-amnestic mild cognitive impairment, or AD.

All participants provided written informed consent. Apolipoprotein E (APOE) genotyping was performed as part of the ADNI and HCP protocols.

### 2.2 PET Biomarker

Aβ-PET imaging was performed using the [^18^F]Florbetapir radiotracer in the ADNI and [^11^C]PiB in the HCP. Standardized uptake value ratios were converted into Centiloid units using a validated methodology.^20^ Aβ-PET positivity was defined as a Centiloid value ≥20.

### 2.3 Plasma p-tau217 assay platforms

Plasma was collected, processed, and stored according to ADNI and HCP protocols: in the ADNI, plasma samples were analyzed using ALZpath (Quanterix)^21^, Janssen p-tau217+ (Quanterix)^22^, Fujirebio Lumipulse^23^, and C2N PrecivityAD2^24^ assays; and HCP sample aliquots using ALZpath, Janssen, Fujirebio, and the University of Pittsburgh in-house assay.^19^ The Fujirebio Lumipulse G p-tau217 assay included in this study is part of the FDA-cleared Lumipulse G pTau217/β-Amyloid 1-42 Plasma Ratio test for the initial evaluation of Alzheimer’s disease.^25^

### 2.4 Harmonization Methods

**Table 1** lists the harmonization methods evaluated in this study, including whether each method incorporates covariate adjustment. Method descriptions and their assumptions are included in **appendix p 2 and 5**, respectively.

**Table 1.**
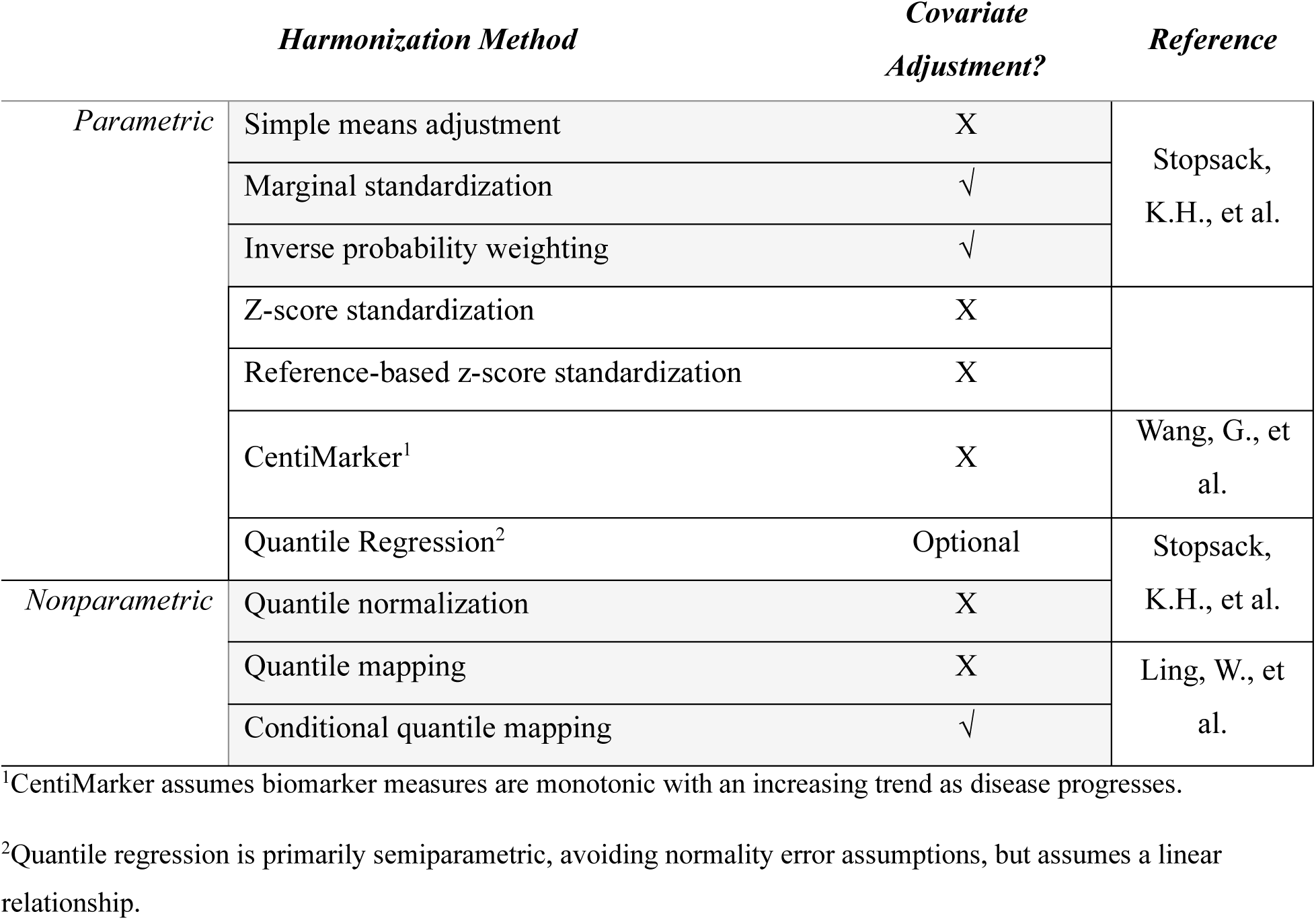
Harmonization methods.

For cohort effects, we considered the following methods: (1) simple means adjustment, (2) marginal standardization, (3) inverse probability weighting (IPW), (4) z-score standardization, (5) reference-based z-score standardization, (6) CentiMarker, (7) quantile regression, (8) quantile normalization, and (9) conditional quantile mapping.

For assay effects, we used only a subset of the methods mentioned in the paragraph above for cohort effects, specifically methods 1, 4-8, and quantile mapping (10).

Simple means adjustment, marginal standardization, IPW, quantile regression, and quantile normalization were applied using the ‘batchtma’ R package.^27^ The code to perform quantile mapping and conditional quantile mapping can be made available.

### 2.5 Evaluation metrics

#### 2.5.1 Evaluation of Technical Variability

Technical variability can be broken down into how it impacts the data distribution across assays or cohorts: differences in mean (additive effects), variance (multiplicative effects), and broader differences in distribution shape (distributional effects).^28^ Additive and multiplicative effects were evaluated by comparing assay-specific or cohort-specific means and SDs, respectively. Empirical distributional differences were evaluated using the k-sample Anderson-Darling test statistic (Ak^2^).^29^ A higher Ak^2^ indicates greater divergence between empirical distributions. These evaluation metrics were applied to both analyses, with further stratification by Aβ-positivity in the cohort-effect analysis. Because cohort differences include both technical and biological variability, we also evaluated residual cohort effects in multiple linear regression models, adjusting for age, sex, race, diagnostic group, APOEε4 status, and Aβ positivity.

Additionally, within-subject variability was quantified in the assay effect analysis using 1-ICC, where ICC is the intraclass correlation coefficient. The ICC was estimated using a linear mixed-effects model (LMM) with assay platform as a fixed effect and a subject-specific random intercept (**appendix p 6**). Higher 1-ICC values indicate greater within-subject disagreement, reflecting assay effects and other residual variation. Bootstrapped 95% confidence intervals (CI) were calculated using 1,000 resamples.^30^

#### 2.5.2 Evaluation of Biological Variability

In the assay effect analysis, we examined the impact of harmonization methods on biological variability by comparing mean differences between CI and CU diagnostic groups estimated using adjusted LMMs (**appendix p 7**).

In the cohort effect analysis, effect sizes between diagnostic groups were estimated with unadjusted Cohen’s d for unequal variances before and after each harmonization method. Adjusted mean differences between diagnostic groups were also estimated using multiple linear regression. Lastly, the predictive performance of p-tau217 for Aβ positivity was evaluated before and after harmonization. The pooled data were randomly split into training (80%) and testing (20%) sets for model development and evaluation. To ensure fair comparison across harmonization methods and assay platforms, the same training and testing sets were used throughout all analyses. Logistic regression models were fit using the pooled training data, with adjustment for age, sex, and APOEε4 carrier status. Model discrimination in the testing set was quantified using the area under the receiver operating curve (AUC), and calibration was assessed using the Brier score.^31^

### 2.6 Statistical Analysis

Descriptive statistics (means, SD) were calculated for all continuous variables. Spearman’s rank correlation coefficients were computed for each pair of assay platforms within the ADNI and HCP cohorts to assess platform concordance. A natural log transformation was applied to p-tau217 values before harmonization to reduce the influence of outliers and improve the validity of parametric harmonization methods.

Assay effects were first evaluated within the ADNI and HCP cohorts using repeated-measures data across assay platforms. Cohort effects were then evaluated by pooling ADNI and HCP data and analyzing each shared assay platform (ALZpath, Fujirebio, and Janssen) separately. Because conditional quantile mapping requires complete-case analysis, individuals with missing data were excluded from the pooled dataset to ensure comparability across harmonization methods. Harmonization methods and evaluation metrics were applied as described in Sections 2.4 and 2.5, respectively. For methods with covariate adjustment, age, sex, race, diagnostic group, APOEε4 status, and Aβ positivity were included as covariates.

All statistical analyses were performed in R version 4.4.1 (2024-06-14).^32^ All statistical tests were two-sided, with statistical significance defined as p < 0.05. No multiple comparison corrections were performed.

## 3 Results

### 3.1 Assay Effects

330 participants from two cohorts (ADNI: n = 219, HCP: n = 111) with repeated measurements of p-tau217 from multiple assay platforms were included in the study (**Table 2**). In ADNI, raw natural log-transformed p-tau217 (ln-ptau217) mean (SD) concentrations ranged from -2.66 (0.596) using the Janssen assay to 1.07 (0.595) using the C2N assay. Similarly, in HCP, ln-ptau217 mean (SD) concentrations ranged from -3.26 (0.575) using Janssen to 2.54 (1.20) using the in-house assay. The ADNI cohort showed high correlations across all assay platforms (Spearman’s r = 0.81-0.89), whereas in HCP, commercially available assay platforms had moderate correlations (Spearman’s r = 0.46-0.66), and the in-house assay had weak correlations (Spearman’s r = 0.37) (appendix p 8).

**Table 2.** Participant demographic and clinical characteristics.

|  | ADNI |  | HCP |  |
| --- | --- | --- | --- | --- |
|  | CU<br>n = 97 | CI<br>n = 122 | CU<br>n = 67 | CI<br>n = 44 |
| **Age, years, mean (SD)** | 81.4 (7.2) | 78.4 (7.8) | 66.8 (8.2) | 65.8 (10.3) |
| **Female, n (%)** | 57 (59%) | 55 (45%) | 43 (64%) | 26 (59%) |
| **Race** |  |  |  |  |
| White | 93 (96%) | 118 (97%) | 44 (66%) | 21 (48%) |
| Non-White | 4 (4.1%) | 4 (3.3%) | 23 (34%) | 23 (52%) |
| **Education, y, mean (SD)** | 16.5 (2.5) | 15.9 (2.8) | NA | NA |
| **APOEε4 carriers, n (%)** | 38 (39%) | 64 (52%) | 19 (28%) | 16 (36%) |
| **Centiloid, mean (SD)** | 35.04 (39.96) | 49.32 (39.57) | 13.10 (25.00) | 24.06 (36.13) |
| **Aβ PET positive, n (%)** | 48 (51%) | 87 (71%) | 12 (24%) | 12 (36%) |
| **pTau-217, pg/mL, mean (SD)** |  |  |  |  |
| ALZpath | 0.49 (0.29) | 0.65 (0.39) | 0.31 (0.15) | 0.46 (0.56) |
| Fujirebio | 0.24 (0.32) | 0.32 (0.29) | 0.10 (0.07) | 0.38 (1.49) |
| Janssen | 0.07 (0.04) | 0.09 (0.06) | 0.04 (0.03) | 0.05 (0.04) |
| C2N | 2.99 (2.18) | 3.97 (2.93) | NA | NA |
| Pittsburgh In-House | NA | NA | 19.35 (20.69) | 33.43 (73.95) |

Abbreviations: NA = not applicable; CI = cognitively impaired; CU = cognitively unimpaired; p-tau217 = phosphorylated tau 217

**Figure 1** shows the impact of each harmonization method on the additive, multiplicative, and distributional assay effects, represented by alignment of assay means, variability, and overall distribution, respectively. **Table 3** summarizes the technical variability metrics evaluated for each method.

**Figure 1.**
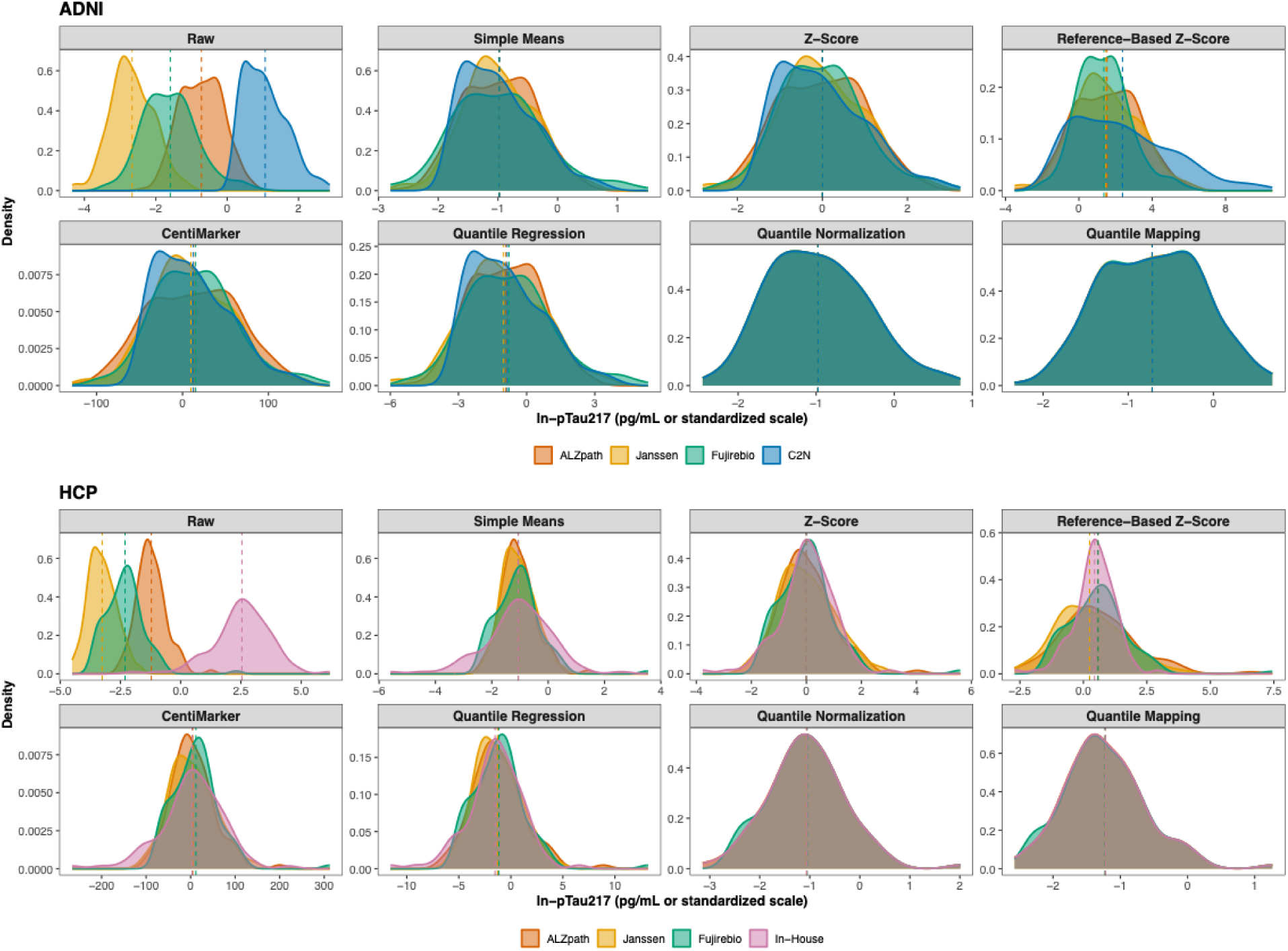
Distributions of ln-ptau217 values across assay platforms before (raw) and after harmonization. Density plots of ln-ptau217 values are shown for each assay platform in the ADNI (top) and HCP (bottom) cohorts before (labelled as Raw) and after application of each harmonization method. Colored dashed vertical lines represent the assay-specific mean. CentiMarker, z-score standardization, and reference-based z-score standardization are unitless. Z-score-based methods represent standard deviation units from the mean. CentiMarker uses a 0-100 scale, with 0 representing normal levels and 100 representing nearly maximally abnormal levels. All other methods express values in pg/mL units. ln-ptau217= natural log-transformed phosphorylated tau 217. ADNI = Alzheimer’s Disease Neuroimaging Initiative. HCP = Human Connectome Project.

**Table 3.**
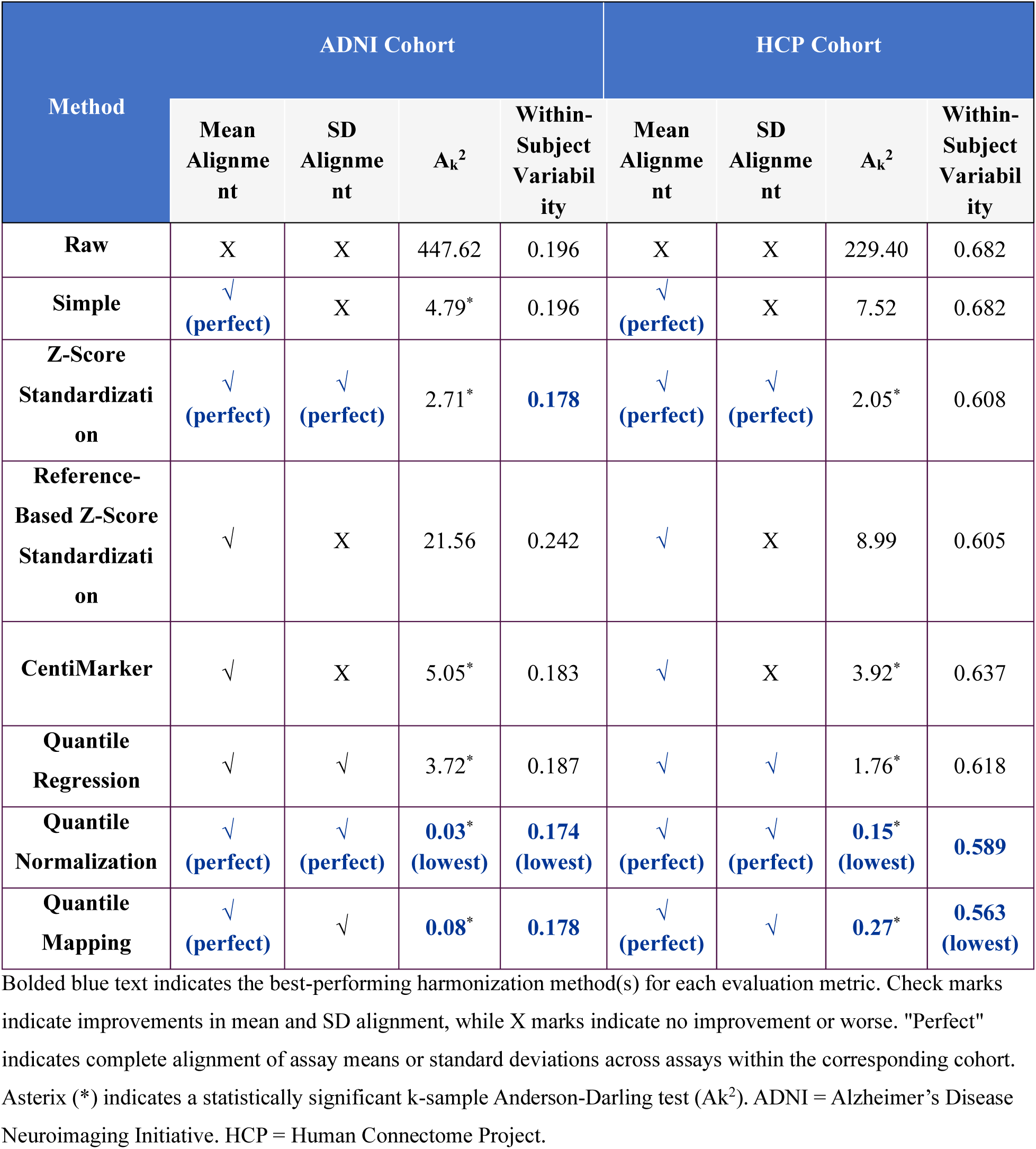
Statistical results for evaluation metrics used to assess assay effects.

| Method | ADNI Cohort |  |  |  | HCP Cohort |  |  |  |
| --- | --- | --- | --- | --- | --- | --- | --- | --- |
| | Mean Alignment | SD Alignment | $A_k^2$ | Within-Subject Variability | Mean Alignment | SD Alignment | $A_k^2$ | Within-Subject Variability |
| Raw | X | X | 447.62 | 0.196 | X | X | 229.40 | 0.682 |
| Simple | ✓<br>(perfect) | X | 4.79* | 0.196 | ✓<br>(perfect) | X | 7.52 | 0.682 |
| Z-Score Standardization | ✓<br>(perfect) | ✓<br>(perfect) | 2.71* | **0.178** | ✓<br>(perfect) | ✓<br>(perfect) | 2.05* | 0.608 |
| Reference-Based Z-Score Standardization | ✓ | X | 21.56 | 0.242 | ✓ | X | 8.99 | 0.605 |
| CentiMarker | ✓ | X | 5.05* | 0.183 | ✓ | X | 3.92* | 0.637 |
| Quantile Regression | ✓ | ✓ | 3.72* | 0.187 | ✓ | ✓ | 1.76* | 0.618 |
| Quantile Normalization | ✓<br>(perfect) | ✓<br>(perfect) | **0.03\***<br>(lowest) | **0.174**<br>(lowest) | ✓<br>(perfect) | ✓<br>(perfect) | **0.15\***<br>(lowest) | **0.589** |
| Quantile Mapping | ✓<br>(perfect) | ✓ | **0.08\*** | **0.178** | ✓<br>(perfect) | ✓ | **0.27\*** | **0.563**<br>(lowest) |

Bolded blue text indicates the best-performing harmonization method(s) for each evaluation metric. Check marks indicate improvements in mean and SD alignment, while X marks indicate no improvement or worse. "Perfect" indicates complete alignment of assay means or standard deviations across assays within the corresponding cohort. Asterisk (\*) indicates a statistically significant k-sample Anderson-Darling test ( $A_k^2$ ). ADNI = Alzheimer's Disease Neuroimaging Initiative. HCP = Human Connectome Project.

All harmonization methods improved assay alignment to varying degrees. Mean (SD) values are listed in **appendix p 9**, the k-sample Anderson-Darling test statistics in **appendix p 10**, and within-subject variability in **appendix p 11-12**. Quantile normalization and quantile mapping were the most effective, producing identical means and variability across assays, eliminating significant distributional differences, and achieving the greatest reductions in within-subject variability in both cohorts. Z-score standardization also produced identical means and variability and reduced within-subject variability, although it was less effective at reducing distributional differences. Simple means adjustment aligned assay means but did not alter variability or within-subject variability. Quantile regression and CentiMarker produced moderate improvements but were less effective than these other methods. Reference-based z-score standardization consistently performed the worst, showing minimal improvement in mean and variability alignment, failing to eliminate significant distributional differences across assays, and increasing within-subject variability in the ADNI cohort.

We further evaluated the impact of data harmonization on downstream biomarker analysis using LMMs, which included assay, diagnostic group, and other demographic covariates (**Figure 2, appendix p 13**). Every harmonization method, except for reference-based z-score standardization in ADNI, removed the significant mean differences observed between each assay platform and ALZpath (reference) prior to data harmonization. Additionally, all harmonization methods preserved the statistically significant difference between CI and CU groups, though the precision of the estimate varied across the methods. Simple mean adjustment yielded estimates identical to those of the raw model in both cohorts. In contrast, quantile regression yielded larger effect estimates and higher standard errors in both cohorts. Notably, quantile mapping reduced the standard error, resulting in narrower confidence intervals and more precise estimates in both cohorts. As a sensitivity analysis, we repeated the analysis using continuous Centiloid values instead of Aβ-positivity groups (**appendix p 14**). The results showed no meaningful changes.

**Figure 2.**
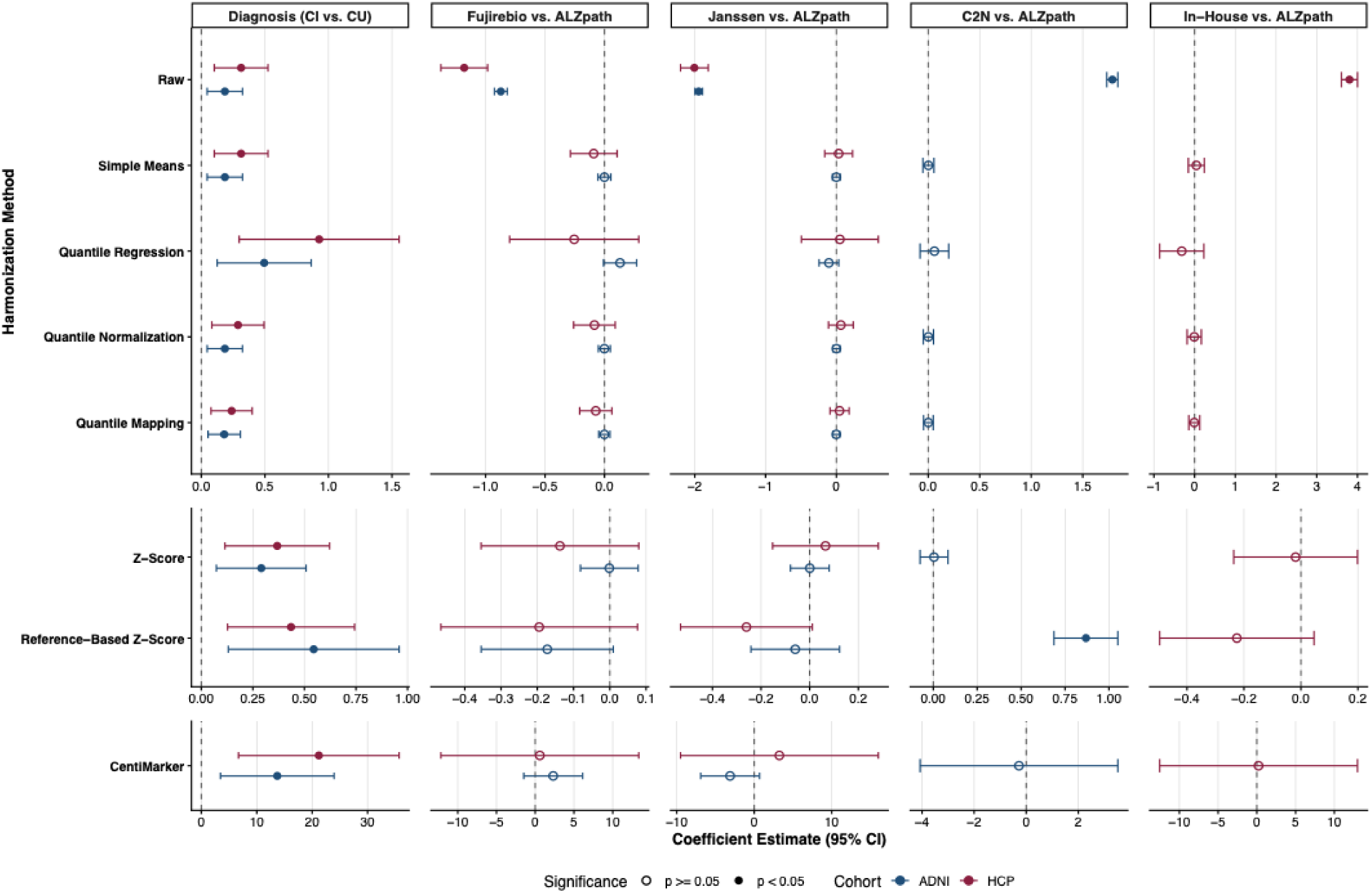
Estimated effects of diagnosis and assay platform before and after harmonization. Linear mixed models were fit for each harmonization method within each ADNI (blue) and HCP (red) cohort. Each model included a subject-specific random intercept and fixed effects for diagnostic group, assay platform, age, sex, race, APOEε4 status, and Aβ-positivity status. The ADNI model additionally adjusted for years of education. Points represent regression coefficients (β) and horizontal lines denote 95% confidence intervals. Filled circles indicate statistically significant associations (p < 0.05), whereas open circles indicate non-significant associations. The diagnosis coefficient represents the adjusted mean difference in ln-ptau217 between CI and CU participants. Assay coefficients represent adjusted mean differences relative to the ALZpath assay (reference). Z-score-based methods express differences in standardized units and CentiMarker expresses differences in CentiMarker units. All other methods remain on the original ln-pTau217 scale. ln-ptau217= natural log-transformed phosphorylated tau 217. ADNI = Alzheimer’s Disease Neuroimaging Initiative. HCP = Human Connectome Project. CI = cognitively impaired. CU = cognitively unimpaired.

### 3.2 Cohort Effects

295 participants pooled from ADNI (n = 212) and HCP (n = 83) were included in the cohort effect analysis (**appendix p 15**). Compared with the HCP, the ADNI cohort was significantly older and had a greater proportion of white, cognitively impaired, and Aβ-positive individuals. Mean p-tau217 concentrations were also significantly higher in ADNI across the ALZpath, Fujirebio, and Janssen assays.

**Appendix p 16-17** shows the impact of data harmonization on overall cohort alignment. Similar to the assay effect analysis, z-score standardization and quantile normalization produced identical empirical distributions across cohorts, simple means adjustment aligned cohort means without changing variability, and reference-based z-score standardization and CentiMarker were the least effective across all assays. Marginal standardization, IPW, quantile regression, and conditional quantile mapping showed moderate improvements.

We further evaluated harmonization methods within Aβ-positive and Aβ-negative groups (**appendix p 18-20**). In the raw data, ADNI consistently had higher ln-ptau217 values than HCP within both Aβ groups across all assays. All harmonization methods preserved Aβ-positive and Aβ-negative group separation but varied in their ability to improve cohort alignment. Conditional quantile mapping produced the greatest improvement in mean, SD, and distributional alignment, followed by marginal standardization, IPW, and quantile regression. In contrast, reference-based z-score standardization and CentiMarker had minimal improvement, whereas simple means adjustment, z-score standardization, and quantile normalization generally worsened cohort alignment within Aβ groups.

Residual cohort effects were then evaluated using multiple linear regression models adjusting for diagnosis and demographic covariates (**Figure 3, appendix p 21**). In the raw data, Fujirebio and Janssen had significant cohort differences, whereas ALZpath did not. Only marginal standardization, IPW, and conditional quantile mapping consistently reduced or kept cohort effects statistically insignificant across all assays. Quantile regression was effective for two assays, whereas reference-based z-score and CentiMarker produced results similar to the raw data. Simple means, quantile normalization, and z-score standardization resulted in significant cohort differences across all assays.

**Figure 3.**
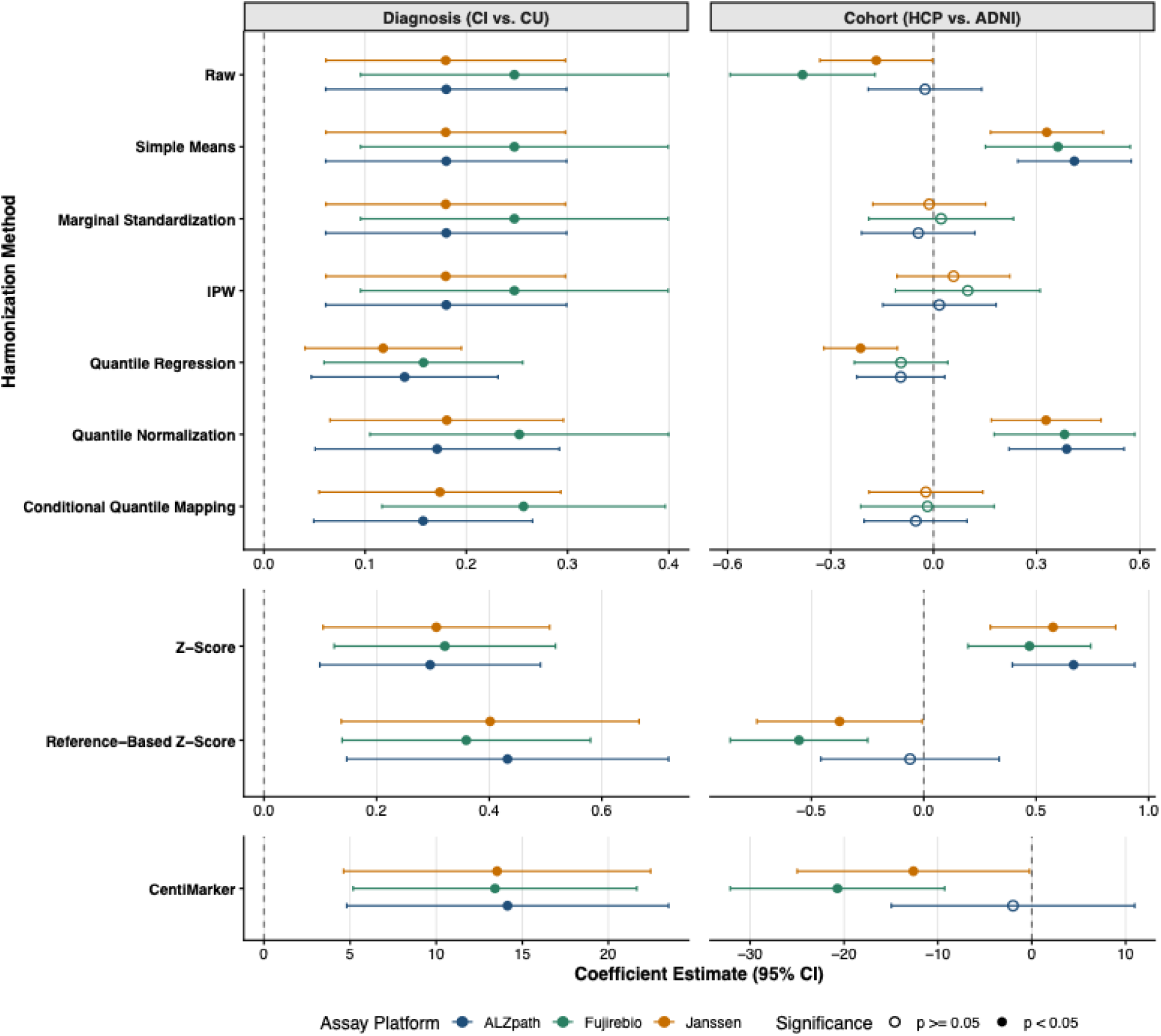
Estimated effects of diagnosis and cohort before and after harmonization. Multiple linear regression models were fit for each harmonization method within each assay: ALzpath (blue), Fujirebio (green), and Janssen (orange). Each model included diagnostic group, cohort, age, sex, race, APOEε4 status, and Aβ-positivity status covariates. The ADNI model additionally adjusted for years of education. Points represent regression coefficients (β) and horizontal lines denote 95% confidence intervals. Filled circles indicate statistically significant associations (p < 0.05), whereas open circles indicate non-significant associations. Z-score-based methods express differences in standardized units and CentiMarker expresses differences in CentiMarker units. All other methods remain on the original ln-pTau217 scale. ln-ptau217= natural log-transformed phosphorylated tau 217. ADNI = Alzheimer’s Disease Neuroimaging Initiative. HCP = Human Connectome Project. CI = cognitively impaired. CU = cognitively unimpaired.

Notably, across all assay platforms, the mean ln-ptau217 difference between the CI and CU groups remained significant. CentiMarker and z-score-based methods yielded coefficients on different scales, limiting direct comparability. Nonetheless, all methods preserved a positive and statistically significant effect. Moreover, while most methods yielded estimates and standard errors similar to those of the raw model, quantile regression and conditional quantile mapping reduced the standard error, producing narrower confidence intervals and greater precision. As a sensitivity analysis, we repeated the analysis using continuous Centiloid values instead of Aβ-positivity groups (**appendix p 22**). The results showed no meaningful changes. Similarly, Cohen’s d estimates showed that all harmonization methods preserved diagnostic group differences with minimal changes (**appendix p 23**).

Lastly, we evaluated the impact of harmonization on downstream predictive performance (**appendix p 24**). Predictive performance and calibration were highly consistent across all harmonization methods. Only conditional quantile mapping improved AUC across all assays, though the gains were modest. Reference-based z-score, CentiMarker, and quantile regression resulted in identical AUC as the raw in all assays.

## 4 Discussion

In this study, we systematically evaluated several harmonization methods to mitigate technical variability in plasma p-tau217, including assay and cohort effects, while preserving biological variability. From these assessments, we found that quantile normalization performed best for combining multi-assay repeated-measures data, whereas conditional quantile mapping performed best for combining pooled multi-cohort data.

In our analysis of repeated-measures data from the ADNI and HCP cohorts, where we expected p-tau217 concentrations to be similar across assays, we observed substantial differences in assay-specific means, SDs, and empirical distributions. Quantile normalization, followed by quantile mapping, was the most effective at reducing these differences while preserving biological variability. On the other hand, reference-based z-score and simple means adjustment performed the worst. In downstream modeling of p-tau217, all methods preserved the significant adjusted mean difference between CI and CU diagnostic groups, while quantile mapping and quantile normalization also improved the precision of the diagnostic group estimate. However, while quantile normalization performed well for repeated measures data across multiple assays, it performed poorly in our analysis of cohort effects.

In our assessment of cohort effects using pooled ADNI and HCP data, we found that conditional quantile mapping, followed by IPW and marginal standardization, performed best, whereas simple means, z-score standardization, and quantile normalization performed worst. Therefore, in this context of pooled data where cohort differences reflect both technical and biological variability, an important distinction emerged between harmonization methods that account for covariate imbalance and those that do not. Methods that explicitly incorporated covariate information, particularly conditional quantile mapping, IPW, and marginal standardization, consistently outperformed other approaches. These methods were most effective at eliminating residual cohort effects in adjusted multiple linear regression models, aligning distributions within Aβ positivity groups, and preserving clinically meaningful differences between diagnostic groups. Together, these results suggest that methods incorporating covariate information are more effective at disentangling technical variation from biological differences.

In contrast, methods that did not account for confounding, including simple means adjustment, quantile normalization, z-score-based approaches, and CentiMarker, were generally less effective. These methods also performed relatively poorly in the assay effects analysis. As expected, simple means adjustment aligned only the assay means while retaining the original variability. Its poor performance suggests that technical variability are not purely additive effects but also involve differences in variance and distributional shape.

Furthermore, reference-based z-score standardization and CentiMarker produced little improvement beyond the raw data, with minimal reduction in technical variability and persistent residual cohort effects. Similarly, although quantile normalization and z-score standardization perfectly aligned marginal distributions across assays and cohorts, significant residual cohort effects remained after covariate adjustment, demonstrating that marginal distribution alignment alone is insufficient to eliminate technical differences. These findings may also reflect a fundamental limitation of z-score-based methods and CentiMarker: rather than preserving the comparable pg/mL numerical scale, they standardize values into assay- or study-specific scales. Consequently, identical standardized values do not necessarily represent the same underlying p-tau217 concentration across assays or cohorts, and downstream analyses should still adjust for cohort or assay to account for these differences in interpretation. In contrast, harmonization methods such as marginal standardization, IPW, and conditional quantile mapping retain measurements on a common scale while reducing technical variation, resulting in more comparable biomarker values across studies.

A few key challenges and limitations should be considered when interpreting these findings. First, although the repeated-measures dataset allowed us to isolate assay effects by holding biological variability constant, we still do not know the true p-tau217 distribution in the absence of technical variability. As a result, harmonization methods were evaluated based on distributional characteristics and the consistency of downstream analyses. Second, the generalizability of these findings may be limited. The harmonization methods we evaluated were applied to a specific set of assay platforms and cohorts. Therefore, their performance may vary depending on study design, population characteristics, and assay-specific differences. Future research should extend these harmonization methods and evaluations to different study designs and additionally use larger datasets. Third, in quantile mapping and conditional quantile mapping, the ALZpath assay and ADNI cohort, respectively, were selected to be the reference. Given that these methods align all distributions to the reference, the choice of reference can influence downstream results, including within-subject variability and effect estimates. Aligning to an assay with lower measurement error and stronger discrimination between CI and CU diagnostic groups may improve comparability and downstream performance, whereas aligning to a noisy assay may carry over that noise to other assays. Future work should evaluate the impact of reference selection on downstream results.

As BBMs become increasingly integrated across assay platforms and studies, systematic evaluation of technical variability and harmonization methods will be essential to improve the reliability and comparability research findings. While efforts to standardize how blood samples are collected, handled, and assayed is critical for quality of downstream data analysis, technical variability is also an inherent part of biomarker studies, especially in larger studies. Therefore, researchers should anticipate for presence of technical variability and have harmonization methods available to mitigate these effects in the data preprocessing step. From our evaluations, we recommend researchers use quantile mapping, or conditional quantile mapping for heterogenous data, to mitigate technical variability in multi-assay or multi-cohort studies. On the other hand, we caution against using z-score based-methods and CentiMarker for the purposes of data harmonization. Overall, our study provides a systematic framework for evaluating technical variability and harmonization methods and offers practical guidance for integrating BBM data across assays or cohorts while preserving meaningful biological signal.

