## Supplementary Materials for "Evaluation of harmonization methods to mitigate assay and cohort effects in plasma p-tau217"

**Supplementary appendix**

**Supplementary Figure 1.** Assay and cohort effect analysis flowchart

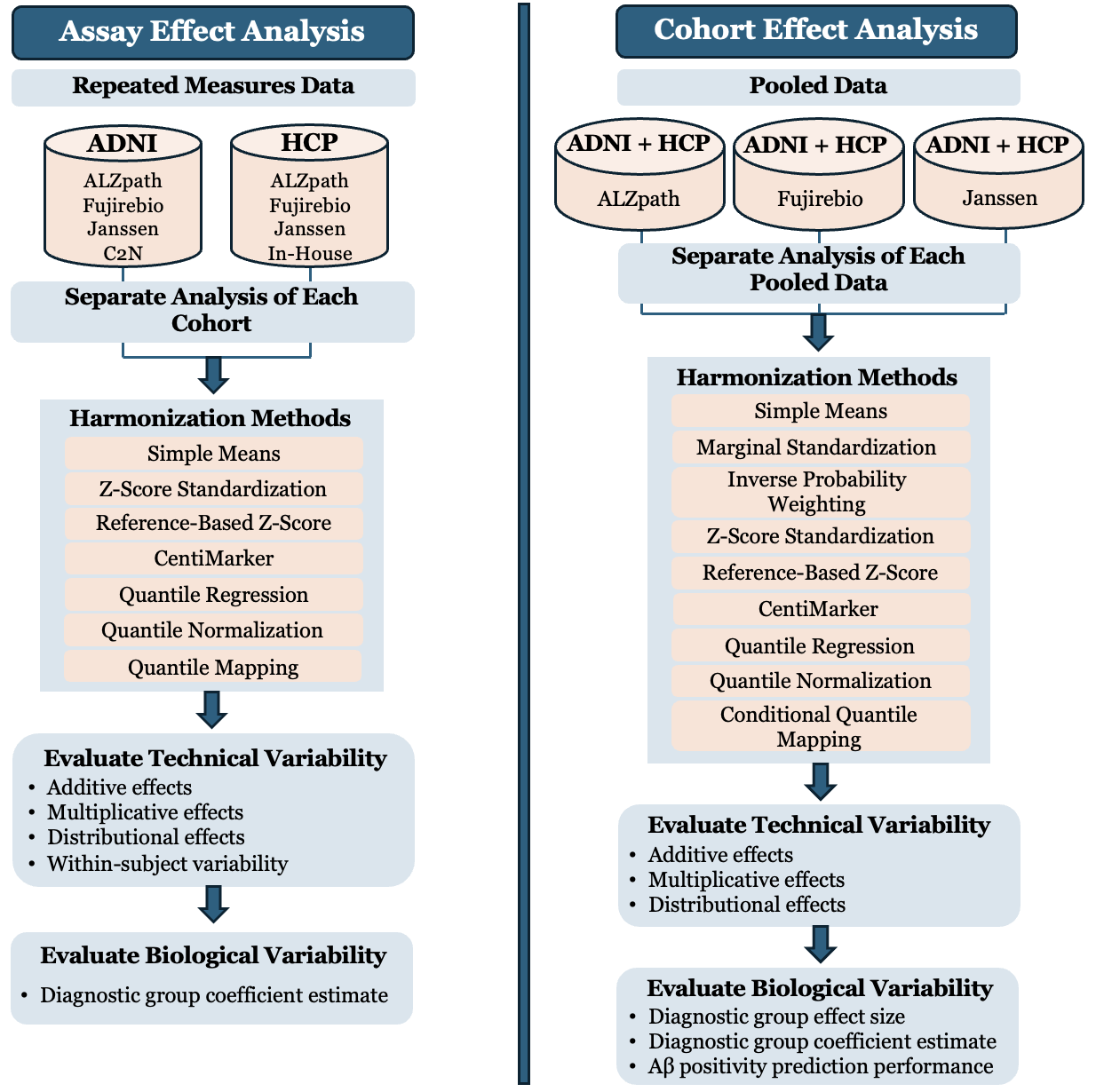

Abbreviations: ADNI = Alzheimer’s Disease Neuroimaging Initiative; HCP = Human Connectome Project

**Harmonization Methods**

1. Simple means adjustment. This method assumes p-tau217 distributions across assays or cohorts have equal means in the absence of technical variability. Technical variability was estimated as the difference in assay- or cohort-specific means, $\bar{y}_{j}$, from the overall mean, $\bar{y}$, where *j* denotes the assay in assay effect analysis or the cohort in the cohort effect analysis. Harmonized values, $\tilde{y}_{ij}$, were obtained by subtracting the estimated technical variability from each observation $y_{ij}$, where *i* indicates participants:

$$\tilde{y}_{ij}={y_{ij}-(\bar{y}}_{j}-\bar{y})$$

2. Marginal standardization. This method extends the simple means adjustment method by incorporating covariate information to account for biological differences between cohorts. It assumes that, in the absence of technical variability, cohorts with similar patient characteristics have equal p-tau217 means. In this study, covariate-adjusted cohort means were estimated using multiple linear regression:

$$y_{ij}=\beta_{0}+\beta_{1}C_{ij}+\gamma^{T}\boldsymbol{X}_{i}+\varepsilon_{ij}$$

where $y_{ij}$ is the raw p-tau217 value for individual i from cohort j and $C_{ij}$ is the cohort indicator, and $\boldsymbol{X}_{i}$ is the vector of covariates representing plausible sources of biological variability between cohorts that should be preserved. Then, harmonized values were obtained by subtracting the estimated adjusted mean difference between cohorts, $\hat{\beta}_{1}$, from raw values.

3. Inverse probability weighting (IPW). This method adjusts for cohort effects by first reweighting observations to balance differences in covariate distributions across cohorts, then aligning the means. First, stabilized weights, ${sw}_{i}$, were calculated for each p-tau217 observation:

$${sw}_{i}=\frac{P\left( C_{i}=j \right)}{P\left( C_{i}=j | \boldsymbol{X}_{\boldsymbol{i}} \right)}$$

where $P(C_{i}=j)$ is the marginal probability of an observation $i$ belonging to their observed cohort $j$. $P(C_{i}=j|\boldsymbol{X}_{\boldsymbol{i}})$ is the probability based on covariates, $\boldsymbol{X}$, estimated using binary (or multinomial) logistic regression:

$$P(C_{i}=j|\boldsymbol{X}_{\boldsymbol{i}})=\frac{1}{1+e^{-(\beta_{0}+\beta_{1}X_{1}+\ldots+\beta_{n}X_{n})}}$$

Stabilized weights were truncated at the 2.5th and 97.5th percentiles to reduce the influence of extreme values. Applying these weights created a pseudo-population in which covariate distributions were balanced across cohorts, thereby reducing confounding. Then, like the marginal standardization method, a weighted marginal linear model was fit to this pseudo-population to estimate cohort-specific mean differences and obtain harmonized values.

4. Z-score standardization. This method centers and scales p-tau217 values within each assay or cohort using assay- or cohort-specific means and standard deviations (SD).
5. Reference-based z-score standardization. This method centers and scales p-tau217 values using the mean and SD of a reference group. For this analysis, the reference group consisted of 15 of the youngest, cognitively unimpaired, APOEε4 negative, and Aβ negative participants across assays/cohorts.

For assay effect analysis, assay-specific means and SDs from the reference group were used to standardize across assay platforms. For cohort effect analysis within the same assay, a common reference mean and SD were estimated from a reference group selected from the pooled ADNI and HCP data. These reference values were then used to standardize p-tau217 across cohorts.

6. CentiMarker. This method standardizes p-tau217 values using two anchor values, CentiMarker-0 and CentiMarker-100, such that values are mapped onto a scale where 0 represents normal levels and 100 represents nearly-maximally abnormal levels. The CentiMarker-0 ($\mu_{CM-0}$) is defined as the mean p-tau217 value among cognitively unimpaired individuals after removing outliers outside the range $Q_{1}-1.5\times IQR$ to $Q_{3}+1.5\times IQR$, where $Q_{1}$ is the 25^th^ percentile, $Q_{3}$ is the 75^th^ percentile, and the interquartile range (IQR) is $Q_{3}-Q_{1}$. The CentiMarker-100 ($\gamma_{CM-100}$) is defined as the 95^th^ percentile of p-tau217 values among cognitively impaired individuals after applying the same outlier exclusion criteria.

For assay effect analysis, assay-specific anchors, $\mu_{CM-0,j}$ and $\gamma_{CM-100,j}$, were used to account for differences in assay scale. The CentiMarker value for each observation $y_{ij}$ was calculated as:

$${CentiMarker}_{ij}=\frac{y_{ij}-\mu_{CM-0,j}}{\gamma_{CM-100,j}-\mu_{CM-0,j}}\times100$$

For cohort effect analysis within the same assay, a common $\mu_{CM-0}$ and $\gamma_{CM-100}$ were derived from pooled data. The CentiMarker value for each observation $y_{ij}$ was calculated as:

$${CentiMarker}_{ij}=\frac{y_{ij}-\mu_{CM-0}}{\gamma_{CM-100}-\mu_{CM-0}}\times100$$

7. Quantile regression. This method assumes that, in the absence of technical variability (i.e. noise), p-tau217 distributions across assays or cohorts with similar characteristics have equal values at select quantiles (e.g., upper and lower quartiles). Assay- or cohort-specific 25th and 75th percentiles were estimated using quantile regression with the Frisch-Newton approach[1]. Harmonized values, $\tilde{y}_{ij}$, were obtained by aligning the 25th percentile and scaling by the interquartile range, following the formula below:

$$\tilde{y}_{ij}=\frac{{(y}_{ij}-\hat{y}_{j}^{\tau=0.25})(\bar{y}^{\tau=0.75}-\bar{y}^{\tau=0.25})}{(\hat{y}_{j}^{\tau=0.75,*}-\hat{y}_{j}^{\tau=0.25,*})}+\bar{y}^{\tau=0.25}-\hat{y}_{j}^{\tau=0.25,*}+\hat{y}_{j}^{\tau=0.25}$$

where $y_{ij}$ indicate the raw p-tau217 value for participant *i* from assay/cohort *j*; $\hat{y}_{j}^{\tau=x}$ the xth quantile of *y* for assay/cohort *j* predicted from unadjusted quantile regression; $\hat{y}_{j}^{\tau=x,*}$the xth quantile of *y* for assay/cohort *j* predicted from adjusted quantile regression; and $\bar{y}^{\tau=x}$the xth quantile of *y* overall. Covariate adjustment wasn’t used in the assay effect analysis.

In assay effect analysis, where covariate adjustment is not needed, the above formula simplifies into:

$$\tilde{y}_{ij}=\frac{{(y}_{ij}-\hat{y}_{j}^{\tau=0.25})}{(\hat{y}_{j}^{\tau=0.75}-\hat{y}_{j}^{\tau=0.25})}(\bar{y}^{\tau=0.75}-\bar{y}^{\tau=0.25})+\bar{y}^{\tau=0.25}$$

8. Quantile normalization. This method assumes the full distribution of p-tau217 values would be identical across assays or cohorts if not affected by technical variability. Values were ranked within each assay/cohort, and values at the same rank were averaged across assays/cohorts. Then, each observation was replaced by the average value corresponding to its rank. For tied values, the average of those values was assigned to their ranks.
9. Quantile mapping. This method aligns the empirical distributions of assays to a reference assay. One assay (ALZpath) was designated as the reference, and its empirical quantiles were estimated using a predefined grid of percentiles, $\tau\in[0,1]$. The same grid was also used to estimate empirical quantiles separately for each non-reference assay based on their observed data. Then, harmonized values were obtained from mapping each non-reference assay observation to the reference scale:

$$\tilde{y}_{ij}=Q_{ref}\left( F_{j}\left( y_{ij} \right) \right)$$

where $F_{j}\left( y_{ij} \right)$ is the percentile of observation $y_{ij}$ in its own assay, and $Q_{ref}(\cdot)$ is the value corresponding to that percentile in the reference assay. As a result, this method preserves each observation’s relative rank while aligning the empirical distribution of the non-reference assay to the reference assay[2]. Linear interpolation was used to ensure a smooth and continuous transformation when estimating percentiles and mapping values between predefined quantile grid points.

10. Conditional quantile mapping. This method extends quantile mapping by incorporating covariate information through quantile regression to estimate percentiles within each cohort[2]. For each observation, its conditional percentile,

$$\tau_{i}=F_{j}\left( y_{ij} | \boldsymbol{X}_{i} \right),$$

was estimated relative to its cohort-specific distribution given covariates, $\boldsymbol{X}_{i}$. This percentile was then mapped to the corresponding value in a reference cohort while holding covariates constant, resulting in an adjusted value on the reference scale:

$$\tilde{y}_{ij}=Q_{ref}\left( \tau_{i}|\boldsymbol{X}_{i} \right)$$

By performing both the percentile estimation and mapping within a covariate-adjusted framework, this approach corrects for cohort effects without removing or distorting covariate-associated variation in the p-tau217 values.

**Harmonization Method Assumptions**

Parametric methods assume that such effects can be described by a specified functional form, typically a linear relationship, and may rely on distributional assumptions such as approximate normality. These methods are sensitive to outliers. In contrast, non-parametric methods do not assume a specific functional or distributional form and instead use data-driven approaches to align p-tau217 distributions across assays or cohorts.

Harmonization methods should be selected based on these assumptions on the data, because it ensures methods work as they’re intended and effectively. Because simple means adjustment relies on mean alignment, it performs best when distributions are approximately symmetric. Marginal standardization and IPW method relies on assumptions of linear regression, including a linear relationship between covariates and raw p-tau217, independence of observations, normality of residuals, and equal variance of residuals (homoscedasticity). Additionally, IPW assumes the cohort assignment model is correctly specified, that all relevant covariates associated with cohort membership and p-tau217 levels are included, and that all individuals have a non-zero probability of belonging to each cohort given their covariates.

For the CentiMarker, researchers recommended that a minimum of 30 observations are used to ensure accurate estimation of the CentiMarker-0 and CentiMarker-100 anchors. Therefore, while anchors can also be derived from additional information, such as APOEε4 negative and Aβ positivity status, only diagnostic status was used in this study to achieve the recommended minimum of 30 data points.

**Model for Quantifying Within-Subject Variability**

The following statistical model was fit to quantify within-subject variability for ADNI.

$$y_{ij}=\beta_{0}+\beta_{1}I(A_{ij}=Janssen)+\beta_{2}I(A_{ij}=Fujirebio)+\beta_{3}I(A_{ij}=C2N)+b_{io}+\varepsilon_{ij}$$

where $y_{ij}$ is the ln-ptau217 level of the i^th^ participant from j^th^ assay (*i* = 1, 2, …, 219, *j* = 1, 2 ,…, 4 (= *n_i_*) since there are *n_i_* = 4 observations per subject); $\beta_{0}$is the mean p-tau217 for the reference assay (ALZpath); I($A_{ij}=k$) is the indicator function equal to 1 if the p-tau217 value was obtained using assay k, and 0 otherwise; $b_{io}$~𝑁(0,$\sigma_{0}^{2}$) is the subject-specific random effect; and $\varepsilon_{ij},$~𝑁(0, $\sigma_{\varepsilon}^{2}$) is the random error term. The terms $\beta_{1}, \beta_{2}, \beta_{3}$represents the mean differences between Janssen, Fujirebio, and C2N and the reference assay (ALZpath), respectively. The model was implemented using the ‘lme4’ package[3].

From this model, the intraclass correlation coefficient (ICC) was used to calculate the 1-ICC, representing the proportion of total variance attributable to within-subject (or between-assay) variability.

$$1-ICC=\frac{\sigma_{\varepsilon}^{2}}{\sigma_{b}^{2}+\sigma_{\varepsilon}^{2}}$$

**Model for Evaluating Biological Variability**

To evaluate the impact of harmonization methods on downstream biomarker analysis, we fit the following random-intercept mixed-effects model for ADNI.

$$y_{ij}=\beta_{0}+\beta_{1}I\left( A_{ij}=Janssen \right)+\beta_{2}I\left( A_{ij}=Fujirebio \right)+\beta_{3}I\left( A_{ij}=C2N \right)+\beta_{4}{Diagnosis}_{i}+\beta_{5}Age_{i}+\beta_{6}Sex_{i}+\beta_{7}Race_{i}+\beta_{8}{A\beta Positivityucation}_{i}+\beta_{9}ApoE{APOE\varepsilon4}_{i}+{\beta_{10}{Education}_{i}+b}_{io}+\varepsilon_{ij}$$

where $y_{ij}$ is the p-tau217 level of participant *i* from assay *j* (*i* = 1, 2, …, 219, *j* = 1, 2 ,…, 4); D_i_ is diagnostic group (ref = cognitively unimpaired); $b_{io}$~𝑁(0,$\sigma_{0}^{2}$) is the subject-specific random effect; and $\varepsilon_{ij},$~𝑁(0, $\sigma_{\varepsilon}^{2}$) is the random error term. The estimate of interest, $\beta_{4}$, represents the adjusted mean difference in p-tau217 concentration between cognitively impaired and cognitively unimpaired groups.

We also fit the model using continuous Centiloid values instead of binary Aβ positivity status to evaluate preservation of variability across Centiloid values. Results for these models using Aβ positivity status and Centiloid are found in appendix p 13 and appendix p 14, respectively.

**Supplementary Figure 2.** Descriptive plots for ADNI and HCP assay platforms

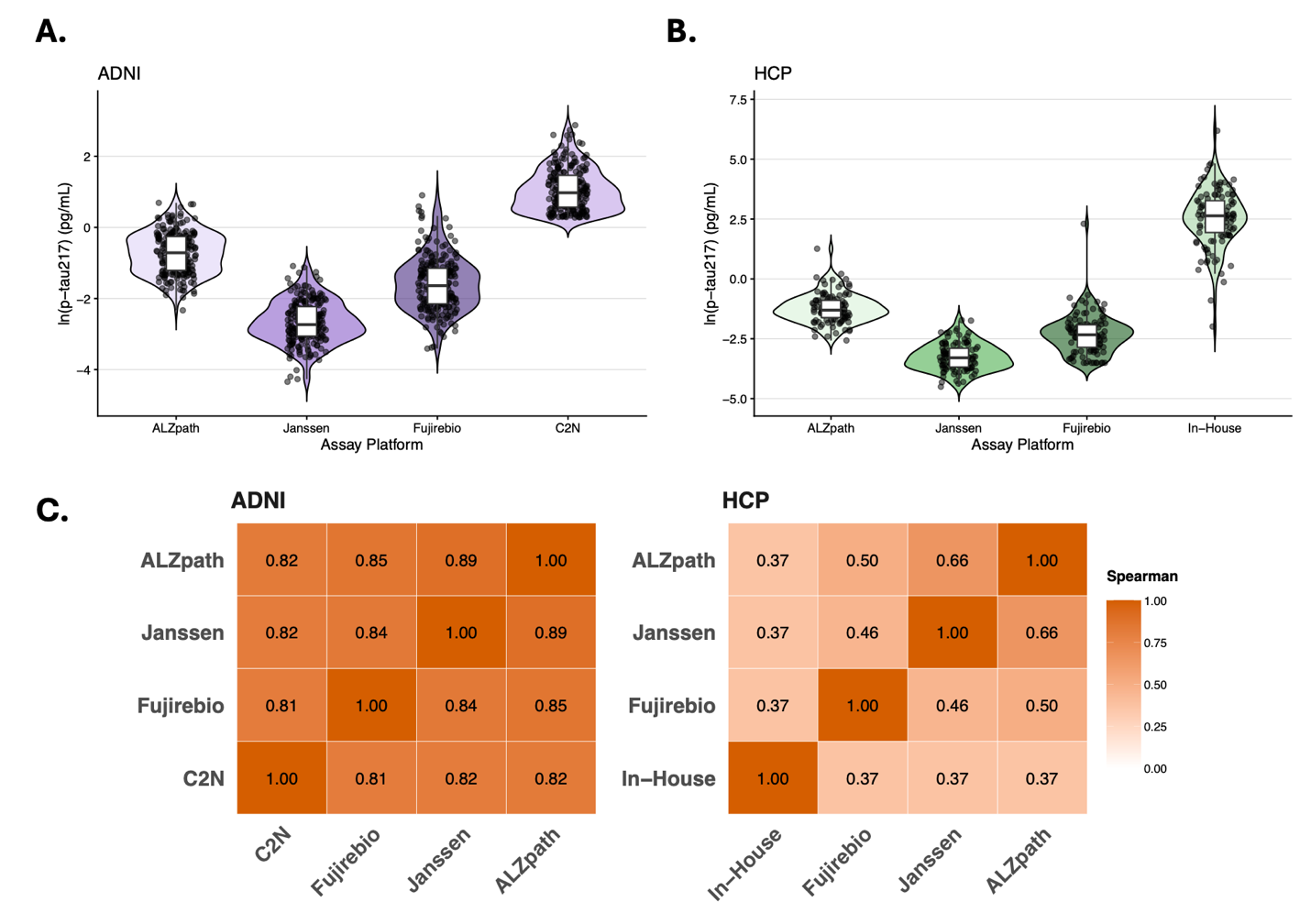

(**A**) Distribution of natural log-transformed plasma p-tau217 concentrations (pg/mL) across four assay platforms in the ADNI cohort. (**B**) Distribution across four assay platforms in the HCP cohort. Violin plots represent the density of values and boxplots indicate median and interquartile range. Substantial differences in absolute concentration ranges across assays are evident in both cohorts. (**C**) Spearman’s rank correlation matrices for pairwise comparisons between assay platforms in ADNI (left) and HCP (right).

**Supplementary Table 1.** ln-pTau217 concentrations before and after harmonization across assay platforms

| Harmonization Method | ADNI Cohort  ln-ptau217 mean (SD), pg/mL | | | | HCP Cohort  ln-ptau217 mean (SD), pg/mL | | | |
| --- | --- | --- | --- | --- | --- | --- | --- | --- |
|  | **ALZpath** | **Janssen** | **Fujirebio** | **C2N** | **ALZpath** | **Janssen** | **Fujirebio** | **In-House** |
| Raw | -0.719  (0.594) | -2.66  (0.596) | -1.59  (0.769) | 1.07  (0.595) | -1.22  (0.616) | -3.26  (0.575) | -2.31  (0.825) | 2.54  (1.20) |
| Simple Means | -0.977 (0.594) | -0.977 (0.596) | -0.977 (0.769) | -0.977 (0.595) | -1.06 (0.616) | -1.06 (0.575) | -1.06 (0.825) | -1.06 (1.20) |
| Z-Score Standardization | 0 (1.00) | 0 (1.00) | 0 (1.00) | 0 (1.00) | 0 (1.00) | 0 (1.00) | 0 (1.00) | 0 (1.00) |
| Reference-Based Z-Score Standardization | 1.53 (1.74) | 1.47 (1.76) | 1.36 (1.44) | 2.38 (2.68) | 0.571 (1.50) | 0.246 (1.31) | 0.575 (1.23) | 0.437 (0.816) |
| CentiMarker | 12.9 (52.0) | 9.76 (45.5) | 15.3 (48.2) | 12.4 (42.4) | 4.31 (48.7) | 4.02 (50.9) | 11.7 (53.8) | 3.32 (71.5) |
| Quantile Regression | -0.909 (1.54) | -1.02 (1.77) | -0.776 (1.89) | -0.855 (1.60) | -1.23 (2.51) | -1.28 (2.13) | -1.14 (2.56) | -1.52 (2.66) |
| Quantile Normalization | -0.977 (0.635) | -0.977 (0.635) | -0.977 (0.635) | -0.977 (0.635) | -1.06 (0.791) | -1.06 (0.791) | -1.05 (0.772) | -1.06 (0.791) |
| Quantile Mapping | -0.719 (0.594) | -0.719 (0.595) | -0.719 (0.594) | -0.719 (0.594) | -1.22 (0.616) | -1.22 (0.616) | -1.21 (0.617) | -1.22 (0.616) |

CentiMarker, z-score standardization, and reference-based z-score standardization are unitless. Z-score-based methods represent standard deviation units from the mean. CentiMarker uses a 0-100 scale, with 0 representing normal levels and 100 representing nearly maximally abnormal levels. ln-ptau217= natural log-transformed phosphorylated tau 217. ADNI = Alzheimer’s Disease Neuroimaging Initiative. HCP = Human Connectome Project.

**Supplementary Table 2.** Distributional differences between assays before and after data harmonization

| Harmonization Method | ADNI Cohort | | HCP Cohort | |
| --- | --- | --- | --- | --- |
|  | **Ak^2^** | **p-value** | **Ak^2^** | **p-value** |
| Raw | 447.62 | < 0.001 | 229.40 | < 0.001 |
| Simple | 4.79 | 0.1 | 7.52 | 0.008 |
| Z-Score Standardization | 2.71 | 0.5 | 2.05 | 0.8 |
| Reference-Based Z-Score Standardization | 21.56 | < 0.001 | 8.99 | 0.002 |
| CentiMarker | 5.05 | 0.08 | 3.92 | 0.2 |
| Quantile Regression | 3.72 | 0.2 | 1.76 | 0.9 |
| Quantile Normalization | 0.03 | 1.0 | 0.15 | 1.0 |
| Quantile Mapping | 0.08 | 1.0 | 0.27 | 1.0 |

Shaded green signifies no significant distributional differences between assay platforms (p > 0.05). Ak^2^ **=** k-sample Anderson-Darling test statistic. ADNI = Alzheimer’s Disease Neuroimaging Initiative. HCP = Human Connectome Project.

**Supplementary Figure 3.** Quantifying within-subject variability before and after harmonization
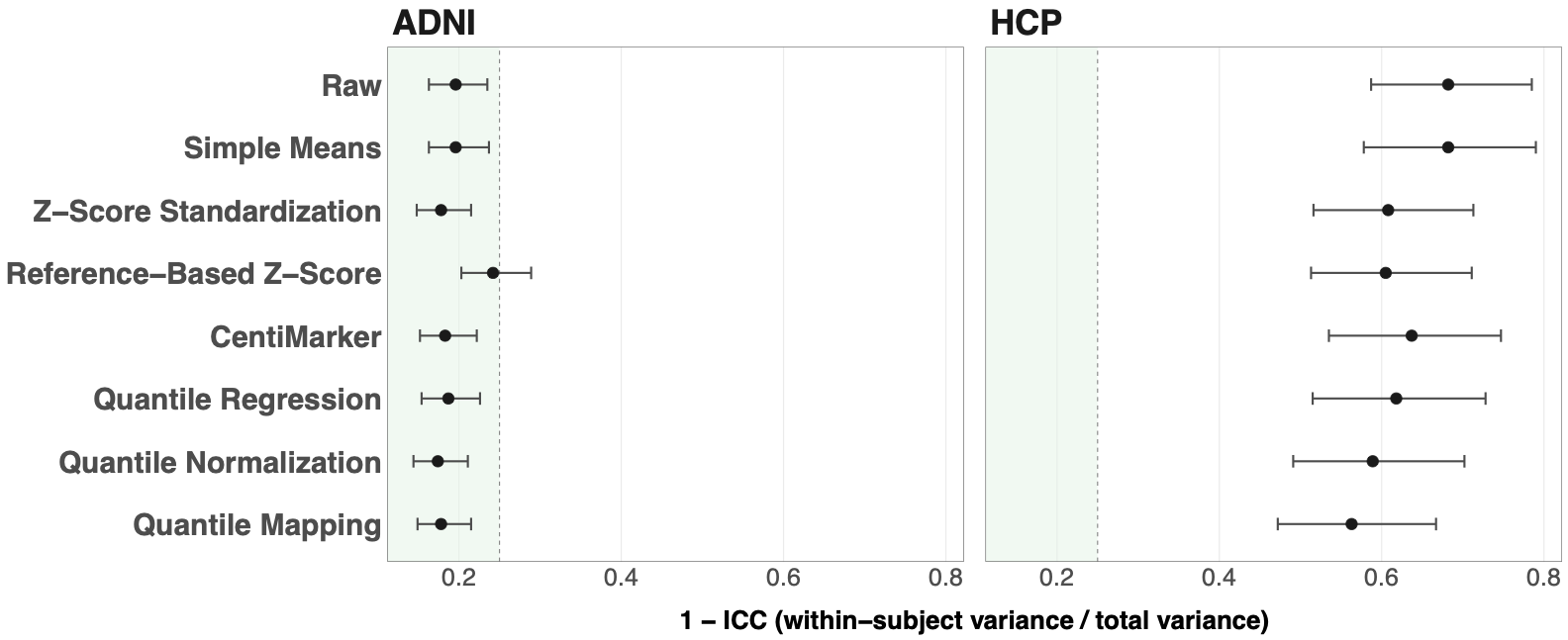

Within-subject variability was quantified as 1-intraclass correlation coefficient (1-ICC), representing the proportion of total variance attributable to within-subject (between-assay) variability. Points indicate 1-ICC estimates with bootstrapped 95% confidence intervals. Reductions in 1–ICC reflect decreased within-subject variability following harmonization in the ADNI (left) and HCP (right) cohorts. A 1-ICC estimate less than 0.25 (green), between 0.25 and 0.5, and greater than 0.5 are indicative of minimal, moderate, and high degrees of within-subject variability, respectively [4]. ADNI = Alzheimer’s Disease Neuroimaging Initiative. HCP = Human Connectome Project.

**Supplementary Table 3.** Within-subject variability before and after harmonization

| Harmonization Method | ADNI Cohort | | HCP Cohort | |
| --- | --- | --- | --- | --- |
|  | **1-ICC** | **95% CI** | **1-ICC** | **95% CI** |
| Raw | 0.196 | (0.163, 0.235) | 0.682 | (0.587, 0.785) |
| Simple Means | 0.196 | (0.163, 0.237) | 0.682 | (0.578, 0.790) |
| Z-Score Standardization | 0.178 | (0.148, 0.215) | 0.608 | (0.516, 0.713) |
| Reference-Based Z-Score Standardization | 0.242 | (0.203, 0.289) | 0.605 | (0.513, 0.711) |
| CentiMarker | 0.183 | (0.152, 0.222) | 0.637 | (0.535, 0.747) |
| Quantile Regression | 0.187 | (0.154, 0.226) | 0.618 | (0.515, 0.728) |
| Quantile Normalization | 0.174 | (0.144, 0.211) | 0.589 | (0.491, 0.702) |
| Quantile Mapping | 0.178 | (0.149, 0.215) | 0.563 | (0.472, 0.667) |

ICC = intraclass correlation coefficient. 1-ICC = within subject variability. ADNI = Alzheimer’s Disease Neuroimaging Initiative. HCP = Human Connectome Project.

Within-subject variability was higher in the HCP cohort than in the ADNI cohort both before and after harmonization. This may reflect differences in cohort composition, as HCP participants are younger and include a greater proportion of CU individuals. At lower biomarker levels, such as those observed in CU individuals, measurement error and assay noise may account for a larger proportion of the observed values, leading to greater relative variability across assays. In contrast, ADNI includes more CI individuals with higher p-tau217 levels, where biological signal may dominate over measurement variability, resulting in lower relative within-subject variability.

**Supplementary Table 4.** Random intercept linear mixed model coefficient estimate for diagnosis

| Harmonization Method | ADNI Cohort | | | HCP Cohort | | |
| --- | --- | --- | --- | --- | --- | --- |
|  | $\boldsymbol{\beta}$ **(SE)** | **95% CI** | **p-value** | $\boldsymbol{\beta}$ **(SE)** | **95% CI** | **p-value** |
| Raw | 0.19 (0.07) | (0.05, 0.32) | 0.01 | 0.31 (0.11) | (0.10, 0.52) | 0.005 |
| Simple Means | 0.19 (0.07) | (0.05, 0.32) | 0.01 | 0.31 (0.11) | (0.10, 0.52) | 0.005 |
| Z-Score Standardization | 0.29 (0.11) | (0.07, 0.51) | 0.01 | 0.37 (0.13) | (0.12, 0.62) | 0.006 |
| Reference-Based Z-Score Standardization | 0.55 (0.21) | (0.13, 0.96) | 0.01 | 0.44 (0.16) | (0.13, 0.74) | 0.007 |
| CentiMarker | 13.70 (5.23) | (3.46, 23.94) | 0.009 | 21.20 (7.39) | (6.71, 35.68) | 0.005 |
| Quantile Regression | 0.49 (0.19) | (0.12, 0.86) | 0.01 | 0.93 (0.32) | (0.30, 1.56) | 0.005 |
| Quantile Normalization | 0.19 (0.07) | (0.05, 0.32) | 0.01 | 0.29 (0.10) | (0.08, 0.49) | 0.007 |
| Quantile Mapping | 0.18 (0.06) | (0.05, 0.31) | 0.006 | 0.24 (0.08) | (0.08, 0.40) | 0.005 |

β coefficient is interpreted as the mean ln-ptau217 (pg/mL) difference between CI and CU holding Aβ positivity, age, sex, race, APOEε4 status, and years of education (ADNI only) constant. For z-score-based methods, coefficient estimates represent adjusted mean differences in standard deviation units. For CentiMarker, coefficient estimates represent adjusted mean differences in CentiMarker units. ADNI = Alzheimer’s Disease Neuroimaging Initiative. HCP = Human Connectome Project.

**Supplementary Table 5.** Random intercept linear mixed model coefficient estimate for diagnosis

| Harmonization Method | ADNI Cohort | | | HCP Cohort | | |
| --- | --- | --- | --- | --- | --- | --- |
|  | $\boldsymbol{\beta}$ **(SE)** | **95% CI** | **p-value** | $\boldsymbol{\beta}$ **(SE)** | **95% CI** | **p-value** |
| Raw | 0.19 (0.06) | (0.06, 0.31) | 0.005 | 0.27 (0.11) | (0.06, 0.48) | 0.01 |
| Simple Means | 0.19 (0.06) | (0.06, 0.31) | 0.005 | 0.27 (0.11) | (0.06, 0.48) | 0.01 |
| Z-Score Standardization | 0.29 (0.10) | (0.09, 0.49) | 0.004 | 0.31 (0.13) | (0.06, 0.56) | 0.02 |
| Reference-Based Z-Score Standardization | 0.54 (0.19) | (0.17, 0.92) | 0.005 | 0.36 (0.15) | (0.06, 0.66) | 0.02 |
| CentiMarker | 13.73 (4.75) | (4.42, 23.04) | 0.004 | 18.14 (7.22) | (3.98, 32.30) | 0.01 |
| Quantile Regression | 0.50 (0.17) | (0.16, 0.83) | 0.004 | 0.79 (0.31) | (0.18, 1.40) | 0.01 |
| Quantile Normalization | 0.19 (0.06) | (0.06, 0.31) | 0.004 | 0.24 (0.10) | (0.04, 0.44) | 0.02 |
| Quantile Mapping | 0.18 (0.06) | (0.07, 0.30) | 0.002 | 0.20 (0.08) | (0.04, 0.35) | 0.01 |

β coefficient is interpreted as the mean ln-ptau217 (pg/mL) difference between CI and CU holding Centiloid, age, sex, race, APOEε4 status, and years of education (ADNI only) constant. For z-score-based methods, coefficient estimates represent adjusted mean differences in standard deviation units. For CentiMarker, coefficient estimates represent adjusted mean differences in CentiMarker units. ADNI = Alzheimer’s Disease Neuroimaging Initiative. HCP = Human Connectome Project.

**Supplementary Table 6.** Participant demographic and clinical characteristics

|  | **Overall** n = 295 | **ADNI** n = 212 | **HCP** n = 83 | **p-value** |
| --- | --- | --- | --- | --- |
| **Diagnosis** |  |  |  | 0.007 |
| CU | 141 (48%) | 91 (43%) | 50 (60%) |  |
| CI | 154 (52%) | 121 (57%) | 33 (40%) |  |
| **Age (years)** | 76.10 (10.01) | 79.79 (7.66) | 66.67 (9.11) | <0.001 |
| **Sex** |  |  |  | 0.2 |
| Female | 156 (53%) | 107 (50%) | 49 (59%) |  |
| Male | 139 (47%) | 105 (50%) | 34 (41%) |  |
| **Race** |  |  |  | <0.001 |
| White | 255 (86%) | 204 (96%) | 51 (61%) |  |
| Non-White | 40 (14%) | 8 (3.8%) | 32 (39%) |  |
| **APOEε4** |  |  |  | 0.051 |
| Without ApoE4 | 169 (57%) | 114 (54%) | 55 (66%) |  |
| With ApoE4 | 126 (43%) | 98 (46%) | 28 (34%) |  |
| **Centiloid** | 36.13 (39.51) | 43.44 (40.38) | 17.46 (30.20) | <0.001 |
| **Aβ positivity** |  |  |  | <0.001 |
| Negative | 138 (47%) | 79 (37%) | 59 (71%) |  |
| Positive | 157 (53%) | 133 (63%) | 24 (29%) |  |
| **ALZpath** | 0.53 (0.38) | 0.58 (0.36) | 0.40 (0.42) | <0.001 |
| **Fujirebio** | 0.24 (0.28) | 0.29 (0.31) | 0.13 (0.11) | <0.001 |
| **Janssen** | 0.07 (0.05) | 0.08 (0.06) | 0.05 (0.03) | <0.001 |

Values are presented as n (%) or mean (SD). P-values were calculated using Pearson's chi-squared test for categorical variables and Welch two-sample t-test for continuous variables.

**Supplementary Figure 4.** Distribution of ln-ptau217 values across cohorts before and after harmonization

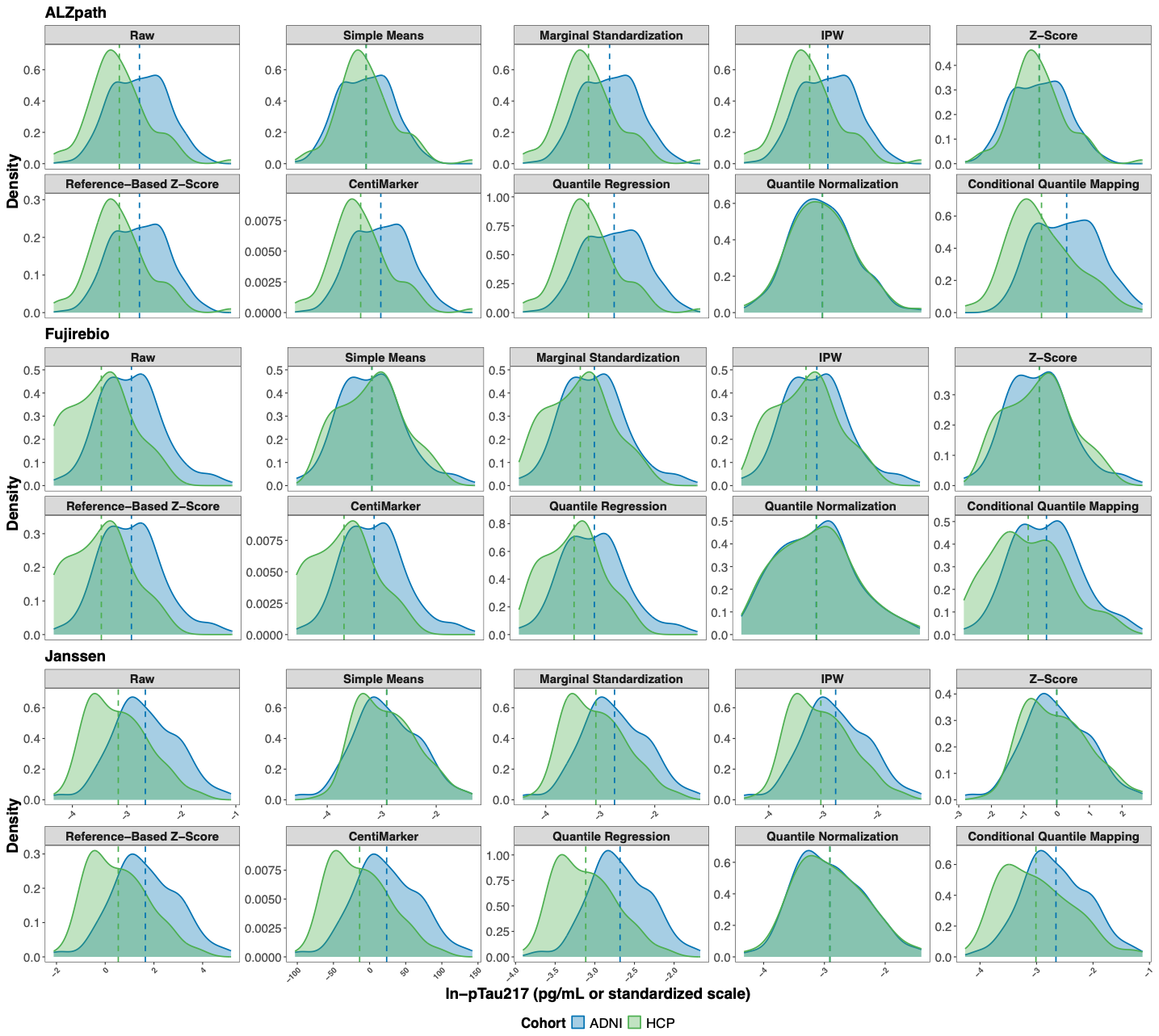

Density plots show the distributions of ln-pTau217 values in the ADNI (blue) and HCP (green) cohorts before and after application of each harmonization method for the ALZpath, Fujirebio, and Janssen assay platforms. Colored dashed vertical lines indicate the cohort-specific mean for each cohort. Z-score standardization, reference-based z-score standardization, and CentiMarker produce standardized values in their own scale. All other methods preserve the pg/mL unit scale. ln-ptau217= natural log-transformed phosphorylated tau 217. ADNI = Alzheimer’s Disease Neuroimaging Initiative. HCP = Human Connectome Project.

**Supplementary Table 7.** ln-ptau217 mean (SD) values across cohorts before and after harmonization

| Harmonization Method | ALZpath | | Fujirebio | | Janssen | |
| --- | --- | --- | --- | --- | --- | --- |
|  | **ADNI** | **HCP** | **ADNI** | **HCP** | **ADNI** | **HCP** |
| Raw | -0.72 (0.59) | -1.15 (0.64) | -1.59 (0.78) | -2.33 (0.76) | -2.66 (0.60) | -3.16 (0.55) |
| Simple | -0.93 (0.59) | -0.93 (0.64) | -1.96 (0.78) | -1.96 (0.76) | -2.91 (0.60) | -2.91 (0.55) |
| Marginal Standardization | -0.71 (0.59) | -1.16 (0.64) | -1.79 (0.78) | -2.13 (0.76) | -2.74 (0.60) | -3.08 (0.55) |
| Inverse-Probability Weighting | -0.74 (0.59) | -1.13 (0.64) | -1.83 (0.78) | -2.09 (0.76) | -2.77 (0.60) | -3.04 (0.55) |
| Z-Score Standardization | 0.00 (1.00) | 0.00 (1.00) | 0.00 (1.00) | 0.00 (1.00) | 0.00 (1.00) | 0.00 (1.00) |
| Reference-Based Z-Score Standardization | 1.79 (1.43) | 0.75 (1.53) | 1.70 (1.13) | 0.62 (1.10) | 1.65 (1.34) | 0.54 (1.24) |
| CentiMarker | 22.11 (46.70) | -12.08 (50.04) | 28.13 (42.22) | -12.16 (41.02) | 23.65 (45.20) | -13.80 (41.65) |
| Quantile Regression | -0.69 (0.47) | -1.10 (0.47) | -1.61 (0.51) | -1.93 (0.45) | -2.68 (0.39) | -3.11 (0.38) |
| Quantile Normalization | -0.94 (0.60) | -0.93 (0.62) | -1.96 (0.76) | -1.95 (0.77) | -2.91 (0.57) | -2.91 (0.58) |
| Conditional Quantile Mapping | -0.71 (0.57) | -1.17 (0.60) | -1.60 (0.72) | -1.99 (0.78) | -2.66 (0.57) | -3.01 (0.63) |

ln-ptau217= natural log-transformed phosphorylated tau 217. ADNI = Alzheimer’s Disease Neuroimaging Initiative. HCP = Human Connectome Project.

**Supplementary Figure 5.** Distribution of ln-ptau217 within Aβ-positivity groups before and after harmonization

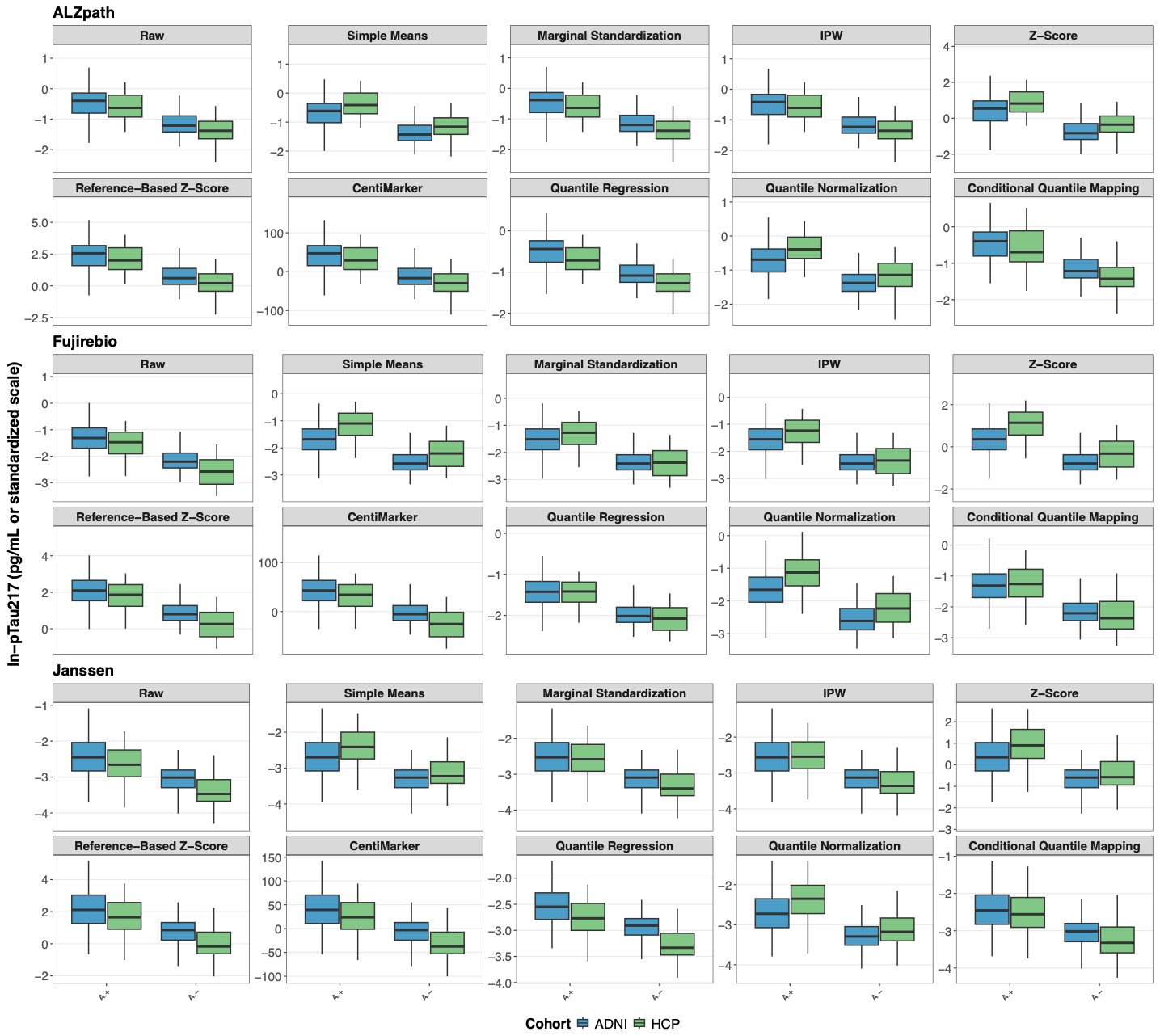

Boxplots show the distribution of ln-pTau217 values within Aβ-positive (Aβ+) and Aβ-negative (Aβ-) groups for the ADNI (blue) and HCP (green) cohorts for each ALZpath, Fujirebio, and Janssen assay platforms. CentiMarker, z-score standardization, and reference-based z-score standardization are unitless. Z-score-based methods represent standard deviation units from the mean. CentiMarker uses a 0-100 scale, with 0 representing normal levels and 100 representing nearly maximally abnormal levels. ln-ptau217= natural log-transformed phosphorylated tau 217. ADNI = Alzheimer’s Disease Neuroimaging Initiative. HCP = Human Connectome Project.

**Supplementary Table 8.** ln-ptau217 mean (SD) values within Aβ-positivity groups before and after harmonization

| Harmonization Method | ALZpath | | | | Fujirebio | | | | Janssen | | | |
| --- | --- | --- | --- | --- | --- | --- | --- | --- | --- | --- | --- | --- |
|  | **A**$\boldsymbol{\beta}$ **Positive** | | **A**$\boldsymbol{\beta}$ **Negative** | | **A**$\boldsymbol{\beta}$ **Positive** | | **A**$\boldsymbol{\beta}$ **Negative** | | **A**$\boldsymbol{\beta}$ **Positive** | | **A**$\boldsymbol{\beta}$ **Negative** | |
|  | **ADNI** | **HCP** | **ADNI** | **HCP** | **ADNI** | **HCP** | **ADNI** | **HCP** | **ADNI** | **HCP** | **ADNI** | **HCP** |
| Raw | -0.46 (0.52) | -0.66 (0.62) | -1.16 (0.42) | -1.35 (0.53) | -1.27 (0.69) | -1.59 (0.65) | -2.13 (0.61) | -2.63 (0.57) | -2.43 (0.55) | -2.65 (0.55) | -3.05 (0.47) | -3.36 (0.40) |
| Simple | -0.67 (0.52) | -0.45 (0.62) | -1.37 (0.42) | -1.13 (0.53) | -1.64 (0.69) | -1.22 (0.65) | -2.50 (0.61) | -2.26 (0.57) | -2.68 (0.55) | -2.40 (0.55) | -3.29 (0.47) | -3.12 (0.40) |
| Marginal Standardization | -0.45 (0.52) | -0.67 (0.62) | -1.15 (0.42) | -1.36 (0.53) | -1.47 (0.69) | -1.39 (0.65) | -2.33 (0.61) | -2.43 (0.57) | -2.51 (0.55) | -2.57 (0.55) | -3.12 (0.47) | -3.29 (0.40) |
| Inverse-Probability Weighting | -0.48 (0.52) | -0.64 (0.62) | -1.18 (0.42) | -1.33 (0.53) | -1.51 (0.69) | -1.35 (0.65) | -2.37 (0.61) | -2.39 (0.57) | -2.54 (0.55) | -2.54 (0.55) | -3.16 (0.47) | -3.25 (0.40) |
| Z-Score Standardization | 0.44 (0.88) | 0.77 (0.97) | -0.74 (0.71) | -0.31 (0.84) | 0.41 (0.89) | 0.98 (0.85) | -0.69 (0.78) | -0.40 (0.75) | 0.38 (0.91) | 0.92 (0.99) | -0.64 (0.79) | -0.38 (0.73) |
| Reference-Based Z-Score Standardization | 2.42 (1.26) | 1.92 (1.48) | 0.73 (1.01) | 0.27 (1.28) | 2.16 (1.00) | 1.69 (0.94) | 0.92 (0.88) | 0.18 (0.83) | 2.16 (1.22) | 1.68 (1.23) | 0.79 (1.06) | 0.07 (0.90) |
| CentiMarker | 42.65 (41.23) | 26.22 (48.31) | -12.49 (33.05) | -27.66 (41.96) | 45.43 (37.40) | 28.03 (35.00) | -1.00 (32.93) | -28.51 (30.88) | 40.90 (41.28) | 24.65 (41.35) | -5.38 (35.80) | -29.43 (30.23) |
| Quantile Regression | -0.49 (0.42) | -0.74 (0.45) | -1.04 (0.33) | -1.25 (0.40) | -1.40 (0.46) | -1.49 (0.39) | -1.96 (0.40) | -2.11 (0.34) | -2.53 (0.35) | -2.76 (0.38) | -2.93 (0.31) | -3.26 (0.28) |
| Quantile Normalization | -0.68 (0.55) | -0.45 (0.58) | -1.36 (0.40) | -1.13 (0.52) | -1.64 (0.66) | -1.19 (0.70) | -2.50 (0.59) | -2.50 (0.59) | -2.69 (0.54) | -2.39 (0.56) | -3.27 (0.41) | -3.12 (0.44) |
| Conditional Quantile Mapping | -0.44 (0.49) | -0.63 (0.58) | -1.16 (0.37) | -1.38 (0.46) | -1.27 (0.63) | -1.25 (0.68) | -2.16 (0.48) | -2.29 (0.60) | -2.43 (0.53) | -2.49 (0.64) | -3.04 (0.42) | -3.22 (0.50) |

CentiMarker, z-score standardization, and reference-based z-score standardization are unitless. Z-score-based methods represent standard deviation units from the mean. CentiMarker uses a 0-100 scale, with 0 representing normal levels and 100 representing nearly maximally abnormal levels. ADNI = Alzheimer’s Disease Neuroimaging Initiative. HCP = Human Connectome Project.

**Supplementary Table 9.** Anderson Darling test statistic comparing ADNI and HCP empirical distributions before and after harmonization

| Harmonization Method | ALZpath | | Fujirebio | | Janssen | |
| --- | --- | --- | --- | --- | --- | --- |
|  | **A**$\boldsymbol{\beta}$ **Positive** | **A**$\boldsymbol{\beta}$ **Negative** | **A**$\boldsymbol{\beta}$ **Positive** | **A**$\boldsymbol{\beta}$ **Negative** | **A**$\boldsymbol{\beta}$ **Positive** | **A**$\boldsymbol{\beta}$ **Negative** |
| Raw | 1.32 | 4.06 | 1.78 | 10.56 | 1.69 | 9.07 |
| Simple | 2.67 | 5.39 | 5.81 | 4.34 | 3.32 | 2.70 |
| Marginal Standardization | 1.63 | 4.80 | 1.02 | 1.96 | 0.48 | 3.39 |
| Inverse-Probability Weighting | 0.97 | 2.77 | 1.72 | 1.76 | 0.39 | 1.77 |
| Z-Score Standardization | 2.04 | 6.37 | 6.14 | 4.21 | 4.02 | 2.14 |
| Reference-Based Z-Score Standardization | 1.32 | 4.06 | 1.78 | 10.56 | 1.69 | 9.07 |
| CentiMarker | 1.32 | 4.06 | 1.78 | 10.56 | 1.69 | 9.07 |
| Quantile Regression | 3.30 | 7.24 | 0.47 | 3.03 | 3.38 | 17.72 |
| Quantile Normalization | 2.95 | 4.86 | 5.15 | 4.31 | 3.92 | 2.60 |
| Conditional Quantile Mapping | 1.69 | 5.77 | 0.37 | 2.26 | 0.56 | 4.45 |

Values represent k-sample Anderson-Darling test statistic. Cells highlighted in green indicate reduced cohort differences compared to the raw. P-values were not evaluated because unlike for the assay effect analysis, we do not expect distributions to become identical and, therefore, are not interested in assessing whether distributions have no significant differences. We are only interested in whether distributional differences are reduced. ADNI = Alzheimer’s Disease Neuroimaging Initiative. HCP = Human Connectome Project.

**Supplementary Table 10.** Multiple linear regression model coefficient estimates for diagnosis

| Harmonization Method | ALZpath | | | Fujirebio | | | Janssen | | |
| --- | --- | --- | --- | --- | --- | --- | --- | --- | --- |
|  | $\boldsymbol{\beta}$ **(SE)** | **95% CI** | $\boldsymbol{\beta}$ **(SE)** | $\boldsymbol{\beta}$ **(SE)** | **95% CI** | **p-value** | $\boldsymbol{\beta}$ **(SE)** | **95% CI** | **p-value** |
| Raw | 0.18 (0.06) | (0.06, 0.30) | 0.003 | 0.25 (0.08) | (0.10, 0.40) | 0.001 | 0.18 (0.06) | (0.01, 0.30) | 0.003 |
| Simple Means | 0.18 (0.06) | (0.06, 0.30) | 0.003 | 0.25 (0.08) | (0.10, 0.40) | 0.001 | 0.18 (0.06) | (0.01, 0.30) | 0.003 |
| Marginal Standardization | 0.18 (0.06) | (0.06, 0.30) | 0.003 | 0.25 (0.08) | (0.10, 0.40) | 0.001 | 0.18 (0.06) | (0.01, 0.30) | 0.003 |
| IPW | 0.18 (0.06) | (0.06, 0.30) | 0.003 | 0.25 (0.08) | (0.10, 0.40) | 0.001 | 0.18 (0.06) | (0.01, 0.30) | 0.003 |
| Z-Score Standardization | 0.30 (0.10) | (0.10, 0.49) | 0.003 | 0.32 (0.10) | (0.12, 0.52) | 0.001 | 0.31 (0.10) | (0.11, 0.51) | 0.003 |
| Reference-Based Z-Score Standardization | 0.43 (0.15) | (0.15, 0.72) | 0.003 | 0.36 (0.11) | (0.14, 0.58) | 0.001 | 0.40 (0.13) | (0.14, 0.67) | 0.003 |
| CentiMarker | 14.15 (4.75) | (4.80, 23.49) | 0.003 | 13.41 (4.18) | (5.18, 21.65) | 0.001 | 13.54 (4.53) | (4.62, 22.46) | 0.003 |
| Quantile Regression | 0.14 (0.05) | (0.05, 0.23) | 0.003 | 0.16 (0.05) | (0.06, 0.26) | 0.002 | 0.12 (0.04) | (0.04, 0.20) | 0.003 |
| Quantile Normalization | 0.17 (0.06) | (0.05, 0.29) | 0.006 | 0.25 (0.07) | (0.10, 0.40) | 0.001 | 0.18 (0.06) | (0.07, 0.30) | 0.002 |
| Conditional Quantile Mapping | 0.16 (0.05) | (0.05, 0.27) | 0.005 | 0.26 (0.07) | (0.12, 0.40) | <0.001 | 0.17 (0.06) | (0.05, 0.29) | 0.004 |

β coefficient is interpreted as the mean ln-ptau217 (pg/mL) difference between CI and CU holding cohort, Aβ positivity, age, sex, and APOEε4 status constant. For z-score-based methods, coefficient estimates represent adjusted mean differences in standard deviation units. For CentiMarker, coefficient estimates represent adjusted mean differences in CentiMarker units.

**Supplementary Table 11.** Multiple linear regression model coefficient estimates for diagnosis

| Harmonization Method | ALZpath | | | Fujirebio | | | Janssen | | |
| --- | --- | --- | --- | --- | --- | --- | --- | --- | --- |
|  | $\boldsymbol{\beta}$ **(SE)** | **95% CI** | $\boldsymbol{\beta}$ **(SE)** | $\boldsymbol{\beta}$ **(SE)** | **95% CI** | **p-value** | $\boldsymbol{\beta}$ **(SE)** | **95% CI** | **p-value** |
| Raw | 0.17 (0.06) | (0.05, 0.27) | 0.003 | 0.24 (0.07) | (0.10, 0.39) | 0.001 | 0.17 (0.06) | (0.06, 0.28) | 0.003 |
| Simple Means | 0.17 (0.06) | (0.06, 0.28) | 0.003 | 0.24 (0.07) | (0.10, 0.39) | 0.001 | 0.17 (0.06) | (0.06, 0.28) | 0.003 |
| Marginal Standardization | 0.17 (0.06) | (0.06, 0.28) | 0.003 | 0.24 (0.07) | (0.10, 0.39) | 0.001 | 0.17 (0.06) | (0.06, 0.28) | 0.003 |
| IPW | 0.17 (0.06) | (0.06, 0.28) | 0.003 | 0.24 (0.07) | (0.10, 0.39) | 0.001 | 0.17 (0.06) | (0.06, 0.28) | 0.003 |
| Z-Score Standardization | 0.28 (0.09) | (0.10, 0.46) | 0.003 | 0.32 (0.10) | (0.13, 0.51) | 0.001 | 0.29 (0.10) | (0.10, 0.48) | 0.003 |
| Reference-Based Z-Score Standardization | 0.41 (0.14) | (0.14, 0.68) | 0.003 | 0.35 (0.11) | (0.14, 0.57) | 0.001 | 0.38 (0.13) | (0.13, 0.62) | 0.003 |
| CentiMarker | 13.45 (4.45) | (4.69, 22.21) | 0.003 | 13.21 (4.00) | (5.31, 21.10) | 0.001 | 12.68 (4.22) | (4.37, 21.00) | 0.003 |
| Quantile Regression | 0.13 (0.04) | (0.05, 0.22) | 0.003 | 0.15 (0.05) | (0.06, 0.25) | 0.001 | 0.11 (0.04) | (0.04, 0.18) | 0.003 |
| Quantile Normalization | 0.16 (0.06) | (0.05, 0.27) | 0.005 | 0.25 (0.07) | (0.11, 0.39) | 0.001 | 0.17 (0.05) | (0.06, 0.28) | 0.002 |
| Conditional Quantile Mapping | 0.15 (0.05) | (0.05, 0.25) | 0.004 | 0.25 (0.07) | (0.12, 0.38) | <0.001 | 0.16 (0.06) | (0.05, 0.27) | 0.004 |

β coefficient is interpreted as the mean ln-ptau217 (pg/mL) difference between CI and CU holding cohort, Centiloid, age, sex, and APOEε4 status constant. For z-score-based methods, coefficient estimates represent adjusted mean differences in standard deviation units. For CentiMarker, coefficient estimates represent adjusted mean differences in CentiMarker units.

**Supplementary Table 12.** Cohen’s d unequal variance test comparing CI with CU

| Harmonization Method | ALZpath | Fujirebio | Janssen |
| --- | --- | --- | --- |
| Raw | 0.53 | 0.60 | 0.55 |
| Simple | 0.45 | 0.51 | 0.52 |
| Marginal Standardization | 0.53 | 0.57 | 0.53 |
| Inverse-Probability Weighting | 0.52 | 0.56 | 0.52 |
| Z-Score Standardization | 0.45 | 0.51 | 0.46 |
| Reference-Based Z-Score Standardization | 0.53 | 0.60 | 0.55 |
| CentiMarker | 0.53 | 0.60 | 0.55 |
| Quantile Regression | 0.54 | 0.58 | 0.56 |
| Quantile Normalization | 0.43 | 0.53 | 0.47 |
| Conditional Quantile Mapping | 0.54 | 0.62 | 0.52 |

**Supplementary Table 13.** Impact of harmonization methods on predictive performance for $A\beta$ positivity

| Harmonization Method | ALZpath | | Fujirebio | | Janssen | |
| --- | --- | --- | --- | --- | --- | --- |
|  | **AUC (95% CI)** | **Brier Score** | **AUC (95% CI)** | **Brier Score** | **AUC (95% CI)** | **Brier Score** |
| Raw | 0.76 (0.63, 0.88) | 0.20 | 0.87 (0.77, 0.96) | 0.15 | 0.74 (0.61, 0.87) | 0.21 |
| Simple | 0.76 (0.63, 0.88) | 0.21 | 0.84 (0.74, 0.95) | 0.16 | 0.76 (0.64, 0.88) | 0.22 |
| Marginal Standardization | 0.76 (0.63, 0.88) | 0.20 | 0.86 (0.76, 0.96) | 0.15 | 0.74 (0.62, 0.87) | 0.21 |
| Inverse-Probability Weighting | 0.76 (0.63, 0.88) | 0.20 | 0.86 (0.76, 0.96) | 0.15 | 0.75 (0.62, 0.87) | 0.22 |
| Z-Score Standardization | 0.76 (0.63, 0.88) | 0.20 | 0.85 (0.75, 0.95) | 0.15 | 0.76 (0.64, 0.88) | 0.21 |
| Reference-Based Z-Score Standardization | 0.76 (0.63, 0.88) | 0.20 | 0.87 (0.77, 0.96) | 0.15 | 0.74 (0.61, 0.87) | 0.21 |
| CentiMarker | 0.76 (0.63, 0.88) | 0.20 | 0.87 (0.77, 0.96) | 0.15 | 0.74 (0.61, 0.87) | 0.21 |
| Quantile Regression | 0.76 (0.64, 0.88) | 0.20 | 0.87 (0.76, 0.96) | 0.15 | 0.74 (0.62, 0.87) | 0.21 |
| Quantile Normalization | 0.75 (0.63, 0.88) | 0.21 | 0.85 (0.74, 0.95) | 0.16 | 0.76 (0.63, 0.88) | 0.21 |
| Conditional Quantile Mapping | 0.79 (0.75, 0.90) | 0.19 | 0.88 (0.79, 0.97) | 0.14 | 0.75 (0.62, 0.88) | 0.21 |

AUC = area under the receiver operating curve.
